# A fast and accurate calculation method for light induced isomerization of retinal proteins in real time

**DOI:** 10.64898/2026.02.27.707937

**Authors:** Philipp Althoff, Kristin Labudda, Udo Höweler, Mathias Lübben, Klaus Gerwert, Carsten Kötting, Till Rudack

**Affiliations:** Center for Protein Diagnostics (PRODI), Biospectroscopy, Ruhr University Bochum, Bochum, Germany; Department of Biophysics, Ruhr University Bochum, Bochum, Germany; Biomolecular Simulations and Theoretical Biophysics Group, Faculty of Biology and Biotechnology, Ruhr University Bochum, Bochum, Germany; CHEOPS, Altenberge, Germany; Structural Bioinformatics Group, Regensburg Center for Biochemistry, Regensburg Center for Ultrafast Nanoscopy, University Regensburg, Regensburg, Germany

**Keywords:** quantum biology, molecular mechanism, molecular dynamics simulations, photo-switchable pro-teins, photocycle, optogenetic tool design

## Abstract

Retinal is a chromophore covalently bound to various photoreceptors. Its photo-induced isomerization triggers a series of structural changes named photocycle, leading to diverse biological functions. Despite tremendous advances in structural biology and artificial intelligence-driven structure prediction, it remains challenging to analyze all photocyclic intermediates. Here, we present an optimized computational approach to calculate RSBH^+^ isomerization and its induced structural changes based on a classical molecular mechanics approach using quantum mechanically improved retinal force field parameters. Isomerization is induced by an excited state restraint which is subsequently relaxed to allow the return to the electronic ground state. We applied this approach to the key protein of optogenetics, Channelrhodopsin-2 from *Chlamydomonas reinhardtii* (*Cr*ChR2). Besides the reformation of the *all-trans/*CN-*anti* ground state, we observed the production of a mixture of two isomeric states 13-*cis*/CN- *anti* and 13-*cis*/CN-*syn*. These findings agree with the previously found branched photocycle model based on experimental data. Our calculations show an asymmetric potential energy landscape of the excited state leading to a corresponding isomerization state distribution. Unlike earlier publications, our procedure describes the retinal photoisomerization on the natural timescale of 500 fs. As our newly derived retinal force field parameter set precisely relies on quantum biological knowledge, it assists to improve the refinement of experimental structure biological data. Our readily customizable strategy provides mechanistic insights at high spatio-temporal resolution, which permits accurate structural predictions of early photocycle intermediates. These insights will stimulate the rational design of optogenetic tools thus providing improved diagnostic and therapeutic approaches for neuronal and other diseases.

**Highlights:**

- universal method to study molecular mechanism of optogenetic tools
- retinal photo-isomerization calculation in real time
- prediction of branched photo cycle agrees with experimental IR spectroscopic results
- detected asymmetric excited state potential energy landscape
- assists to improve structural model refinement of retinal proteins

**Graphical Abstract:** 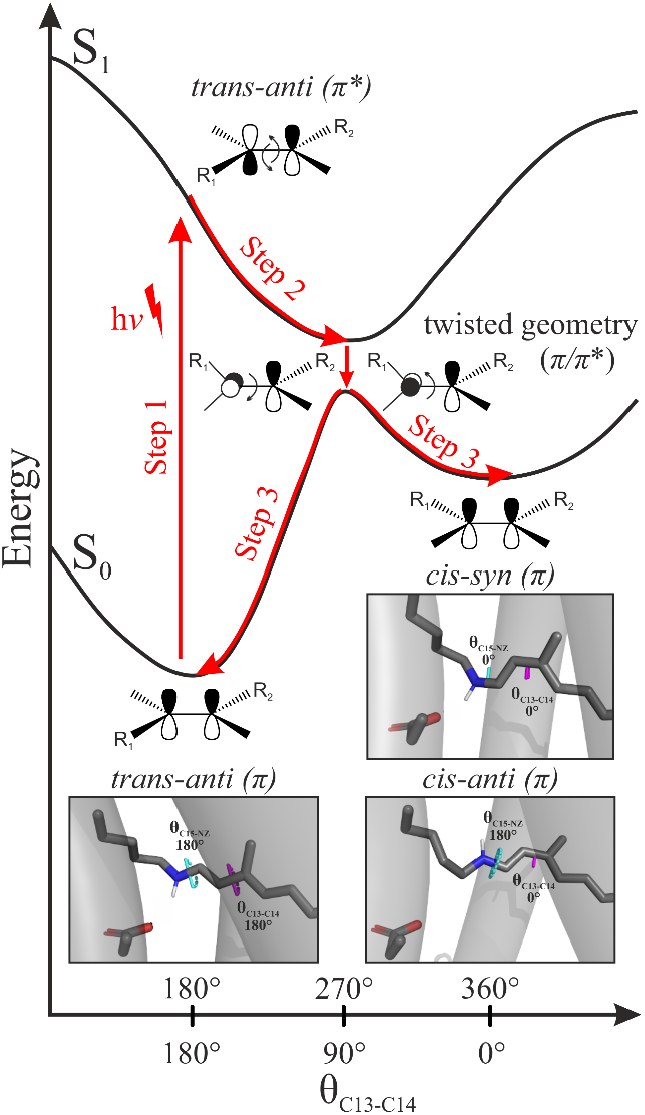

## 1. Introduction

### 1.1 Retinal Proteins

Retinal proteins are light-sensitive molecules that play crucial roles in the triggering of cellular functions, notably in photo-responsive cells [1]. These photoreceptor molecules control important physiological functions e.g. by serving as membrane channels, ion pumps, or G protein coupled receptor (GPCR) activated signaling cascades [2–4]. The ability of photoreceptors to undergo conformational changes upon light absorption makes them pivotal to the field of optogenetics, in which light is used to control biological processes [3, 5–9]. In optogenetics, light-activatable proteins (such as channelrhodopsins) are introduced into tissues through genetic manipulations. These proteins allow precise control of cellular functions, such as neuron activation, by using light [10–13]. It is a powerful tool used to manipulate specific cells within tissues, often for studying neural circuits and behavior [4, 8, 9, 14–18]. Understanding the molecular mechanisms of retinal proteins is essential to the design of novel proteins with tailor-made optogenetic features for use in specific applications [19–21]. Combination of results from infrared (IR) spectroscopic approaches [22–24] with recent advances in 4D structural biology, such as XFEL [25–27] and time-resolved X-ray crystallography [28–30], alongside with computational tools like artificial intelligence driven structure prediction[31–33] and molecular dynamics (MD) simulations[34–36], offers critical insights into structure and dynamics of retinal proteins.

Retinal is the essential cofactor triggering cell physiological effects such as vision and phototaxis [2]. In form of type I and type II rhodopsins it occurs within all domains of life [37–41]. While the exact stimulus transmission varies greatly, all rhodopsins share the common feature to control protein activity by the molecular configuration of the bound retinal [2, 11, 42, 43]. The retinal chromophore is an aldehyde derivative of vitamin A [37–41]. Instead of existing freely, in rhodopsins this cofactor is usually covalently attached to the apoprotein: The retinal carbonyl is coupled to the amino function of a conserved, membrane integrally located lysine to form a protonatable imine group, which is named protonated Schiff-base (RSBH^+^).

### 1.2 Photo-Isomerization

Key feature of the RSBH^+^ as a chromophore is the conjugated *π*-electron-system, including the doublebond between the nitrogen (atom name NZ) and its neighboring carbon atom of the retinal (C15) (see Figure S1) and additional five single- and five double carbon-carbon-bonds. The delocalization of *π*-electrons in conjugated systems lowers the energy-gap between the highest occupied molecular orbital (HOMO) and the lowest unoccupied molecular orbital (LUMO). Free retinal exhibits an absorption maximum of 370 nm, while the maximum of the RSBH^+^ varies depending on the protein-environment around ∼ 500 nm. The photoexcitation S0 → S1 induces a change in the bonding pattern from bonding to antibonding character between carbon atoms C13 and C14 thus favoring a 90° or 270° minimum for the passing θ_C13-C14_ torsion (Figure S2). The geometry is expected to equilibrate within femtoseconds around one of these new minima. At these minima geometries, the potential energy surfaces of S0 and S1 are extremely close and form an avoided conical intersection. Passing through this energy gap the molecule falls back into the ground state and the double bond character is recovered. Depending on the actual excited state θ_C13-C14_ value on passing, the system falls back to its original position or to the opposite direction. In the second case, this bond is isomerized. The isomerization of retinal is with 50-500 fs known to be one of the fastest reactions in biology [44][45]. The photocycle describes the cascade of conformational changes of the photoreceptor triggered by RSBH^+^ isomerization.

Here, we investigate the impact of RSBH^+^ isomerization within the photocycle of Channelrhodopsin-2 from the green algae *Chlamydomonas reinhardtii* (*Cr*ChR2). *Cr*ChR2 is a light-gated cation channel with high specificity for protons and sodium ions, making it a critical tool in the field of optogenetics [12, 46–49]. In this protein the Schiff base is formed by retinal binding to the conserved lysine residue (K257) located within the seventh transmembrane helix. In the ground state, retinal is in the *all-trans* CN(15)-*anti* configuration, referred to as *trans-anti*, but undergoes isomerization to a 13-*cis* configuration upon light activation [13]. The isomerization is the key mechanistic event that triggers a series of conformational and internal protonation changes in the gate residues of the channel, in particular regarding E90, D253, and E123 [50]. These changes are crucial for the ion conductivity of the channel. Recently two distinct photocycle pathways, resulting in the well and the poorly conducting channel were experimentally identified by combination of spectroscopy and electrophysiology [13]. The branched photocycle model suggests that after the *all-trans* to 13-*cis* isomerization, the C15-NZ bond (see Figure S1) adopts either of the two configurations: 13-*cis* CN(15)-*anti* configuration (*cis-anti*) with the intermediate P_500_(K), leading to the well-conducting channel, or 13-*cis* CN(15)-*syn* (*cis-syn*), resulting in the poorly conducting channel associated with the intermediate P_480_ [13].

### 1.3 Earlier Approaches to Calculate the Dynamics of Photo-isomerization

First attempts to theoretically describe the RSBH^+^ isomerization leading to conformational changes of the photoreceptor rhodopsin using a molecular dynamics approach were made by Arieh Warshel in 1976 [51] using semi-empirical methods for the excitation calculation in combination with molecular mechanics (MM) simulations to describe the retinal movement. He proposed the “bicycle-pedal” model to explain the molecular dynamics of retinal during photoisomerization in rhodopsin. His model showed how the restrictive active site of rhodopsin directs the isomerization of 11-*cis* retinal to an *all-trans* configuration. The process occurs via a concerted rotation of double bonds, ensuring efficiency and specificity. The model explained the ultra-fast nature of the reaction that was at that time thought to be ∼6 ps and the high quantum efficiency by highlighting the protein role in constraining retinal motion to a unique pathway. Klaus Schulten probed three different MM methods to induce isomerization in bacteriorhodopsin [52]. One method used an adiabatic shift of equilibrium torsion angles over 10 ps to guide the retinal to its new configuration gradually. A second method mimicked photoexcitation by instantaneously altering the potential energy surface to destabilize the *trans-anti* state and stabilize the *cis* states.

A third method introduced stepwise changes in torsion angles, allowing structural adjustments to be visualized at each stage. Since that, different quantum mechanical (QM), MM, and hybrid QM/MM approaches were performed to investigate the isomerization of retinal in divers’ proteins. Sen et al. [53] reviewed these isomerization mechanisms of retinal proteins. Using hybrid QM/MM simulations, they explore various proposed mechanisms, including the one-bond-flip (OBF), bicycle-pedal (BP), and hula-twist (HT) models. The study highlights how protein environments optimize the efficiency, speed, and selectivity of RSBH^+^ isomerization. Advanced QM/MM methods, particularly multiconfigurational approaches, have improved our understanding of these mechanisms. The authors emphasize the role of conical intersections in excited-state dynamics and predict that ultrafast X-ray crystallography will soon validate computational models.

Recently, calculations have been also applied to investigate isomerization of retinal in Channelrhodopsin-2 using different approaches to initiate the isomerization process. Ardevol and Hummer initiated their simulations by selecting twelve conformations from the D470 ensemble (representing the dark-adapted state) and using the C12-C13-C14-C15 torsion angle (θ_C13-C14_) as the collective variable to drive the isomerization [54]. The metadynamics method, implemented in PLUMED 2.0 [55], was used to bias this torsion angle. This approach forced the transition to the *cis*-*anti* retinal configuration, which corresponds to the channel-opening state, while avoiding the unwanted *cis*-*syn* configuration via a bicycle movement using a smoothed-wall potential. The bias was updated every 4 ps, with a Gaussian-like potential of 0.15 rad width and 1.2 kJ/mol height. Kuhne et al. [13, 56] and Xin et al. [13, 56] both used a stepwise dihedral rotation approach to simulate the RSBH^+^ isomerization. They rotated the C13-C14 bond in 20° increments to gradually shift from the *all-trans* to 13-*cis* configuration. After each step, they performed 10 ns MD simulations to relax the structure and allow the protein to adjust to the new retinal configuration. This iterative process provided detailed insights into the conformational changes in the retinal binding pocket. Xin et al. only rotated around the C13-C14 bond and thus reached the *cis*-*anti* form. Kuhne et al. extended this approach by also simulating the rotation of the C=N bond, capturing the isomerization to both the *cis-anti* and the *cis*-*syn* form. The role of conformational heterogeneity and protonation equilibria in shaping the photocycle branching of Channelrhodopsin-2, particularly emphasizing the early P480 intermediate and the deprotonation dynamics of E90, was further explored by Bellucci et al. using MM simulations and free energy calculations [57]. Bellucci et al used published experimental information to model the early photo cycle intermediates whereas Kuhne et al. and Xin et al. actually calculated the isomerization reaction. However, the latter two used restraints or freeze groups to also guide the isomerization process by experimental information and enforced either the formation of the *cis*-*anti* configuration or the *cis*-*syn* configuration. Within the method used by Ardevol and Hummer, the isomerization occurred over a span of approximately 80 ps, whereas in Kuhne et al. and Xin et al. this process is extended to over 90 ns. Neither of these approaches captures the natural timescale of RSBH^+^ isomerization, which occurs in around 500 fs. Failure to capture the correct time frame in MD simulations gives the protein environment more time to adapt to the conformational changes of the retina than it naturally has. These unnatural equilibration times allow artificial protein conformations to be obtained.

In this work, we therefore introduce a novel method to induce isomerization spontaneously without bias and simulate the process of RSBH^+^ isomerization within the biological time scale of 500 fs. To allow a natural progression of the isomerization process we induce isomerization by the introduction of an excited state force-field. By that, θ_C13-C14_ is forced towards a 90° and 270° angle. Subsequently the groundstate parameter set was introduced again to allow for spontaneous formation of either an isomerization state or reformation of the ground state.

## 2. Results and Discussion

Key to perform reliable RSBH^+^ isomerization calculations is a well equilibrated ground state structure as starting point. To equilibrate the *Cr*ChR2 dimer structural model to its physiological embedding we performed classical molecular mechanics (MM) simulations. Critical prerequisites for reliable MM simulations are the quality of the starting structure and the used force field parameters; therefore, we checked available experimentally resolved structural models of retinal configurations covalently bound to photoreceptors and the respective retinal topologies. As we noticed inconsistencies between the retinal description and quantum chemically (QC) derived knowledge we optimized the topologies accordingly (see 2.1). Using these optimized topologies, we performed MM simulations of *Cr*ChR2 dimers, embedded in a membrane consisting of the phospholipid 1-palmitoyl-2-oleoyl-sn-glycero-3-phosphocholine (POPC) surrounded by water molecules and sodium and chloride ions at physiological concentrations (see 2.2). We then used structural snapshots from the equilibrated part of the simulation trajectory to initiate RSBH^+^ isomerization calculations using our newly developed approach, considering the natural time scale of the isomerization process (see 2.3).

### 2.1 Retinal parameter optimization

A review of bond length values of retinal configurations bound to photoreceptors within structural models deposited in the Protein Data Bank (PDB) revealed inconsistencies (see Figure 1). The different PDB ligand topologies of retinal often fail to represent the chemically accurate structure and correspondingly the photochemical behavior of retinal. The key issue is that there is no topology for a retinal covalently bound to lysine via a protonated Schiff base that forms RSBH^+^. There are only topologies that describe bound retinal via a non-protonated Schiff base (RSB). However, the usual retinal configuration in the ground state of the photoreceptor is RSBH^+^. QC calculations detailed in Figure S3 underline the clear structural difference between RSB and RSBH^+^.

**Figure 1:**
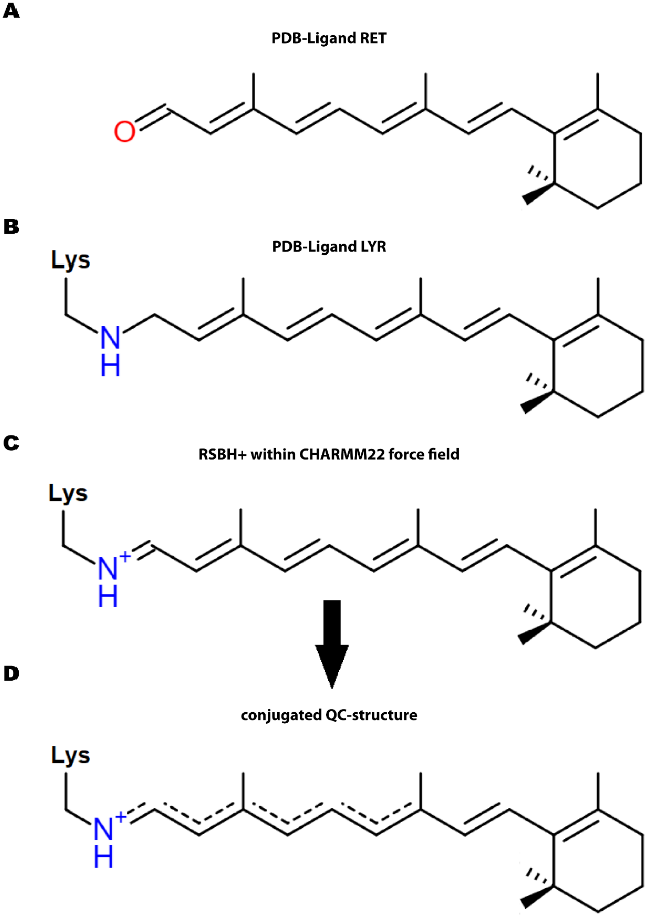
Comparison of PDB-ligand-topologies with quantum chemically derived structures. **A** PDB-ligand RET as unbound retinal-molecule with clearly pronounced bondlength alteration. **B** PDB-ligand LYR, representing retinal bound to lysin but with a chemically incorrect structural formula: C15 is sp^3^ hybridized instead of sp^2^ and has no double bond to the nitrogen, thus no protonated Schiff base is formed. **C** RSBH^+^ represented within the CHARMM22 force field [58]. **D** The RSBH^+^-structure resulting from quantum chemical calculations (Table S6) indicates a conjugated p-system that develops equilibrated bond length from the nitrogen atom to the second methyl-group followed by bond length alternation of the carbon-carbon bonds.

The ligand labeled RET (see Figure 1A) deposited in the PDB ligand data base reflects the isolated retinal molecule chemically correct but does not describe the correct binding character of the RSBH+. Even though it is used as topology for protein bound retinal structures. The absence of any covalent bond to the protein is inconsistent with the optical and biological properties of the complex. The PDB ligand labeled LYR (see Figure 1B) includes the covalent bond between retinal and lysine, but it lacks the critical double bond between the NZ and C15 atoms (see Figure S1) because these atoms have a false sp3 hybridization instead of the chemically correct sp2-hybridization. This wrong assignment leads to disruption of the end of the conjugated p-system, which is an indispensable condition for light absorption, and the RSBH+ isomerization, and the initiation of subsequent photobiological processes.

There exist improved versions of RSBH^+^ in the CHARMM22 force field [58] and in the Amber force field [54] (see Table S1). While these representations reflect the proper double bond between C15 and NZ (see Figure 1C), they feature strictly alternating single- and double bonds instead of equilibrated bond lengths along the conjugated *π*-system. This alternation is not in accordance with QC calculations for RSBH^+^. Table S2 summarizes the results from density functional theory methods and Møller Plesset (MP)2 both yielding highly equilibrated bond length up to the bond C9-C10 carrying the second methylgroup. Our QC calculations (see Table S2) yield a pronounced alternating pattern for the C9-C8 (long), C8-C7 (short), C7-C6 (long), and C6-C5 (short) bonds (see Figure 1D) with bond lengths closer to the 1,3-butadiene values of 1.35 Å and 1.46 Å for a slightly perturbed double bond and formal single bond, respectively. We set the parameter for the equilibrated bond length to 1.4 Å (as in fully conjugated benzene [59]) which is the average of the almost identical bond length results from the advanced QC methods (DFT and MP2). We transferred all RSBH^+^ valence parameters (see Methods Section) to the new set of covalently bound RSBH^+^ parameters to the Optimized Potentials for Liquid Simulations – All Atom (OPLS/AA) force field (see Tables S3-S6). We anticipate that adapting the PDB retinal topologies by incorporating the correct quantum chemical description will improve the experimentally resolved structures of retinal proteins.

### 2.2 Ground-state Simulations

Employing our improved retinal parameters, we conducted three replicates of 1 μs MM simulations of a *Cr*ChR2 dimer, referred to as run1, run2, and run3. Simulation details are provided in the Methods section. The root mean square deviation (RMSD) analysis shown in Figure S4 revealed that all three simulations are ground state equilibrated after 500 ns. Therefore, the subsequent analyses were performed on the last 500 ns of the trajectories. As three replica runs were performed, and the simulation system contained a dimer of *Cr*ChR2 the properties of a total of six retinal binding pockets were analyzed.

First, we analyzed the dihedral angle, defined by the four atoms C12-C13-C14-C15 and the rotation around the C13-C14 axis, referred to as θ_C13-C14_. As an example, Figure 2A shows this dihedral analysis for the simulation trajectory of monomer A from run 3. The average of all equilibrated simulation trajectories resulted in a θ_C13-C14_ value of 194°. We attribute the slight deviation of this value from the theoretically anticipated 180° to steric interactions of retinal located in its protein binding pocket. In addition, we analyzed the evolution of the retinal contact pattern over the simulation time (see Figure 2B). We identified the key interactions that stabilize retinal in the binding pocket: The retinal ring and chain remain largely fixed within the protein-binding pocket by stabilizing van der Waals interactions. The RSBH^+^ of the retinal consistently forms hydrogen bonds with residues in the binding pocket. Figure 2C illustrates the three most prominent hydrogen bond acceptors for the protonated Schiff base, i. e. primarily D253, followed by E123, and by a water molecule that is coordinated by both E123 and D253. Figure S5 gives an overview of the contact pattern of all here performed simulations. We note that due to the restricted amount of simulation time and simulation replica the here obtained distribution percentage of the Schiff base interaction partners should not be overinterpreted. However, it is a justified assumption that all three geometries really occur in nature.

**Figure 2:**
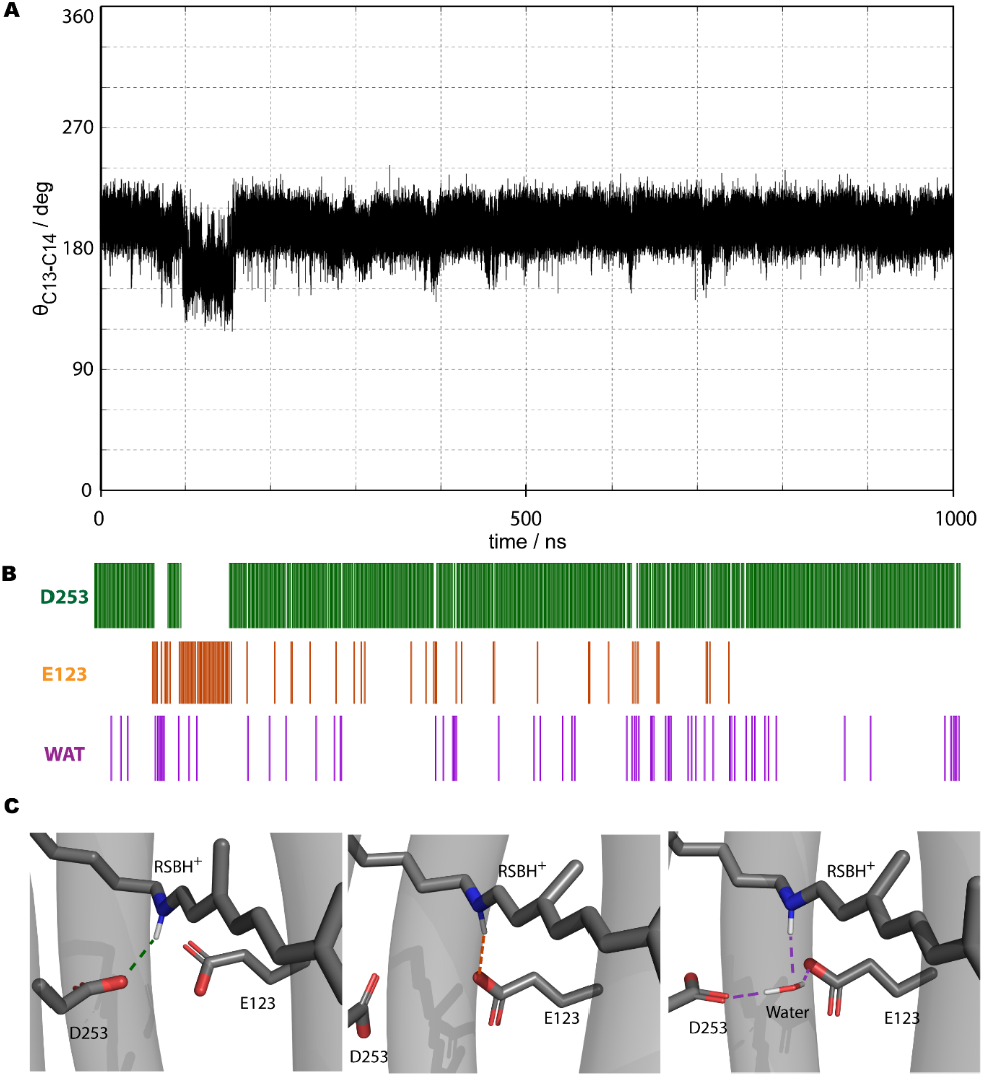
Analysis of geometrical properties of a representative ground state MM simulation. **A** Time response of the dihedral θ_C13-C14_ for 1 ms simulation trajectory of monomer A from run 3. **B** Contact-matrix analysis for the nitrogen atom of the protonated Schiff base to the respective acceptors D253, E123, or a bound water molecule. Bars indicate contacts within the observed time frame of the simulation. **C** Representative structural models of the identified key geometries of the retinal binding pocket occurring during the simulation. The three deduced protein conformations do appear, but their exact relative distribution is unclear due to sampling limitations.

### 2.3 Isomerization calculation strategy

To provide an overview of our applied workflow, Figure 3 illustrates the approach to explore the conformational dynamics and isomerization pathways of *Cr*ChR2. Our strategy is divided into 3 steps using a combination of free and biased classical MM simulations.

**Figure 3:**
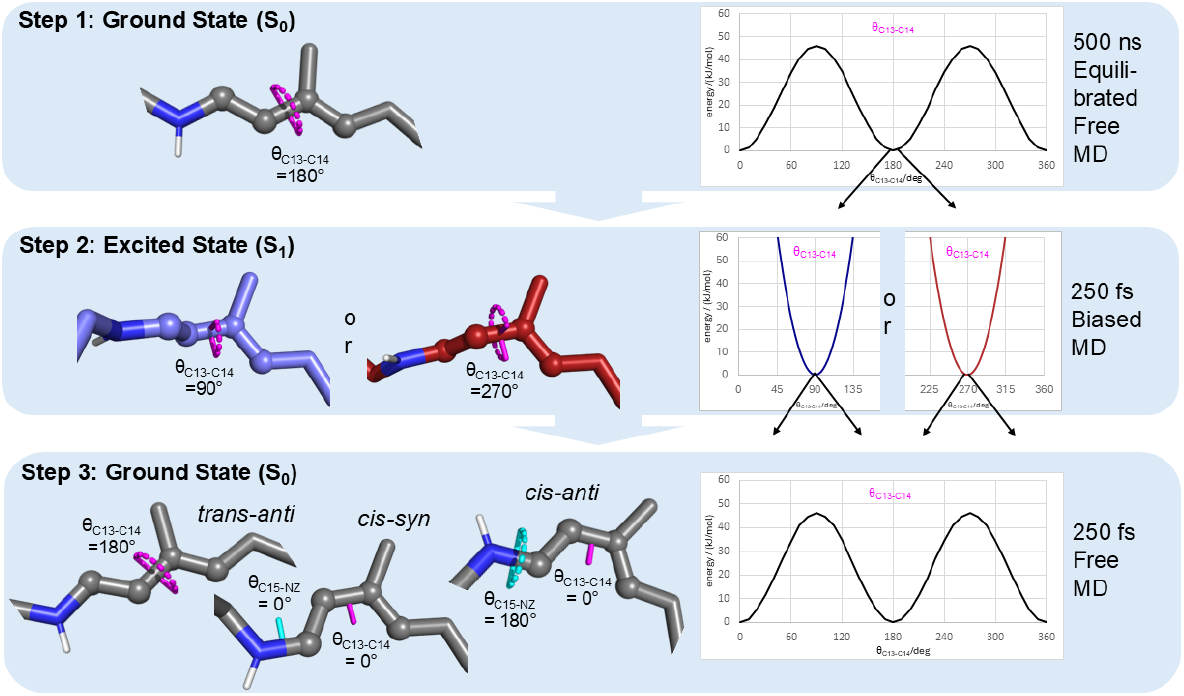
Retinal photo-isomerization workflow. **Step 1:** MM simulations of *Cr*ChR2 in the ground state with the RSBH^+^ initially in all-*trans* CN-*anti* configuration (*trans-anti*, left) and the standard ground-state torsion potential for θ_C13-C14_ (right) were performed. After at least 500 ns equilibration of the MM simulation geometry snapshots were taken every 50 ns to initiate the next step. **Step 2:** A photon (h*v*) induces with equal probability a rotational rotation of θ_C13-C14_ (magenta) towards either 90° or 270°. To account for the light induced changes in retinal configuration, two individual 250 fs MM simulations were performed for each of the snapshots obtained in step 1 with the θ_C13-C14_ excitation state restraint potential (right) at either 90° (blue) or 270° (red) in replacement of the θ_C13-C14_ ground state potential. **Step 3:** To mimic the relaxation to the ground state the simulation from step 2 is extended for another 250 fs after removal of the θ_C13-C14_ excitation state restraint and reintroduction of the θ_C13-C14_ ground state potential (right). This return leads to three unbiased relaxed configurations (left): the *trans*-*anti* ground state, the isomerized 13-*cis* CN-*syn* (*cis-syn*) configuration, and the isomerized 13-*cis* CN-*anti* (*cis-anti*).

**Step1**: The system is equilibrated by an unbiased MM simulation using the standard ground-state torsion potential for θ_C13-C14_. 11 snapshots every 50 ns from the equilibrated last 500 ns of the simulation are selected as starting points for further analysis. **Step 2**: To investigate the photon (h*ν*) induced retinal rotation within the isomerization process, biased 250 fs MM simulations are performed by applying an excited state restraint to θ_C13-C14_. These excited state restraints are introduced to force the system toward the two independent specific rotational minima at either 90° or 270° expected in the first excited state. Here, we probed five different force constants (100, 125, 150, 175, and 200 kJ/mol). Accompanying we set the ground-state torsion potential reflecting θ_C13-C14_ to zero. This procedure allows for controlled exploration of both rotational directions independently within two MM simulations initiated from the same starting structure assuming the same likelihood for both directions. At each of the 11 snapshots from step 1 we performed 20 MM runs with randomly varying velocity distributions of 250 fs duration with the excitation restrain θ_C13-C14_ set to 90° and 20 MM runs with θ_C13-C14_ restrained to 270° leading to in total 440 excitation calculations for one retinal binding pocket and one force constant initiated by one MM run. **Step3**: The ground-state torsion potential is reintroduced, allowing the retinal geometries to relax back to their energy minima during a 250 fs MM simulation. Relaxation toward θ_C13-C14_ of 180° restores the *trans-anti* configuration, while relaxation to 0° results in the isomerized *cis-anti* configuration. Additionally, during this relaxation phase, coupled rotations of the dihedral angle defined by the atoms C14-C15-NZ-CE and the rotation around the NZ-C15 axis, referred to as θ_NZ-C15_, are occasionally occurring thus producing the double-isomerized *cis-syn* configuration. Due to the short simulation time, these geometries are not regarded as equilibrated around a global minimum but rather as localized close to a local minimum that mimics the first intermediate of the photocycle.

Within **step 1** we performed here three independent MM simulations of a dimer resulting in a total of 33 geometries (3 runs times 11 geometries) of which each has two retinal binding pockets for further analysis. The MM runs initiate in **step 2** in total 6,600 excitation runs (33 geometries from step 1 times 20 excitation runs times 2 directions times 5 force constants). After **step 3** there are in total 13,200 (2 times 6,600) protein embedded retinal geometries available for analysis, as each of the resulting geometries is a dimer with two retinal binding pockets.

### 2.4 Isomerization calculation results

To capture diverse configurations of the retinal-binding pocket and ensure accurate representation of system dynamics, we selected eleven geometries from each of the equilibrated simulation trajectory of step 1 to initiate isomerization calculations. Figure 4 represents the results of the three steps of our isomerization strategy here shown for monomer A within simulation run 3. Table S7 shows the details of the resulting data from our isomerization calculation steps. Even though in most cases the retinal remains stable within the simulations of all three steps due to van der Waals interactions, the lysine side chain (K257) including the RSBH^+^ is more flexible, because it adopts different conformations as demonstrated by our retinal contact analysis. The fact of different distinct RSBH+ positions, notable already in the ground state (see Figure 2), enables dynamical adjustment of the chromophore.

**Figure 4:**
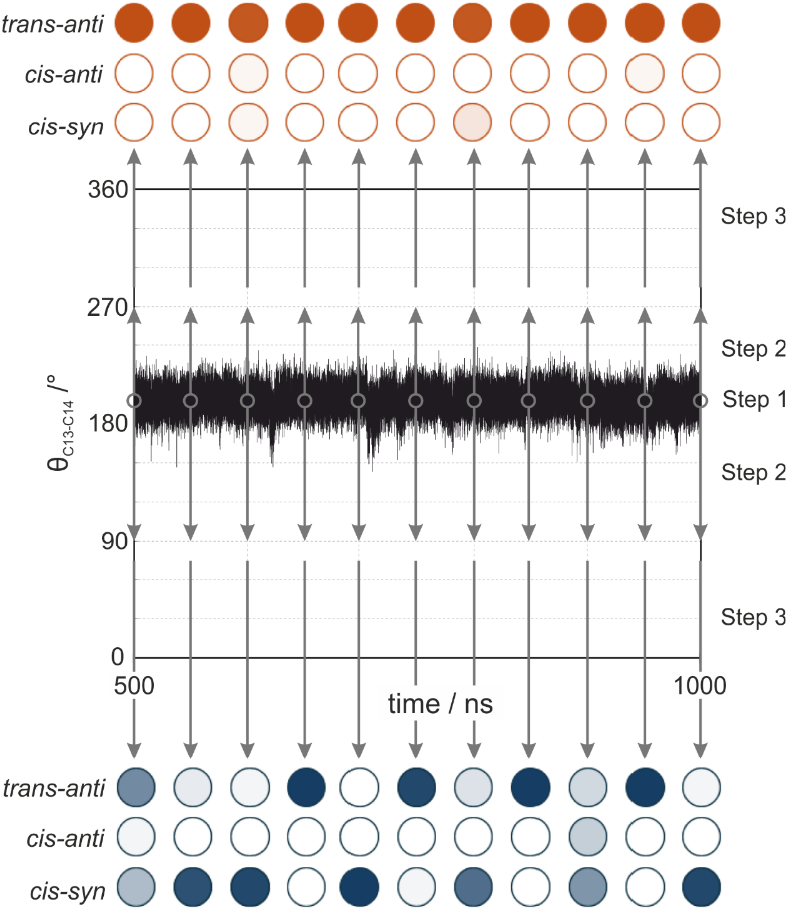
Retinal configurations obtained from exemplary isomerization calculations. Gray circles represent the 11 time points (each 50 ns after at least 500 ns of ground state equilibration), at which step 2 was initiated by taking the corresponding geometry snapshots in two independent calculations, using a restraint for θ_C13-C14_ of either 270° (red) or 90° (blue) was used, followed by subsequent relaxation to the ground state (step 3). The resulting distribution of retinal configurations is represented by the color intensity of the circles, which semi-quantitatively reflects the populations of each configuration. Detailed values are shown in Table S5.

Although the distortion due to electronic excitation was largely localized at θ_C13-C14_, a rotation of θ_NZ-_ C15 from 180° to 0° is also possible. Therefore, we observed three different retinal configurations after isomerization namely trans-anti, cis-syn, and cis-anti (see Figure 3).

To identify the proper force constant adjusted as excited state restraint potential of θ_C13-C14_ that approximates best the resulting retinal distortion after excitation, we tested three different force constants. Figure S6 shows the resulting overall excited state distribution after step 2, represented by the final θ_C13-C14_ angle restrained either towards 90° or 270° obtained after 250 fs of MM simulations. The force constant of 200 kJ/mol proved to be the most reasonable, yielding excited-state geometries with θ_C13-C14_ angles closest to theoretical excited state energy minima. Specifically, geometries subjected to the 90° excited state restraint were distributed around 101°, while those subjected to the 270° excited state restraint were distributed around 263°. We attribute these deviations from the anticipated theoretical values, that are in the same range as observed for the ground state MM simulations, to interactions of the retinal with its protein binding pocket.

The retinal configuration distributions (Figure 5A) of the ground (step 1) and excited state (step 2) yield central points of the potential energy landscape (Figure 5B). Based on the observed 14° shift of the equilibrated ground state minimum observed for retinal bound in *Cr*ChR2 we assume that also the maxima are symmetrically shifted by 14°, even though we do not explicitly calculate them. Our data of the excited state imply an asymmetric distribution of the excited state minima of -7° from the ideal value 270° and +11° from 90°.

**Figure 5:**
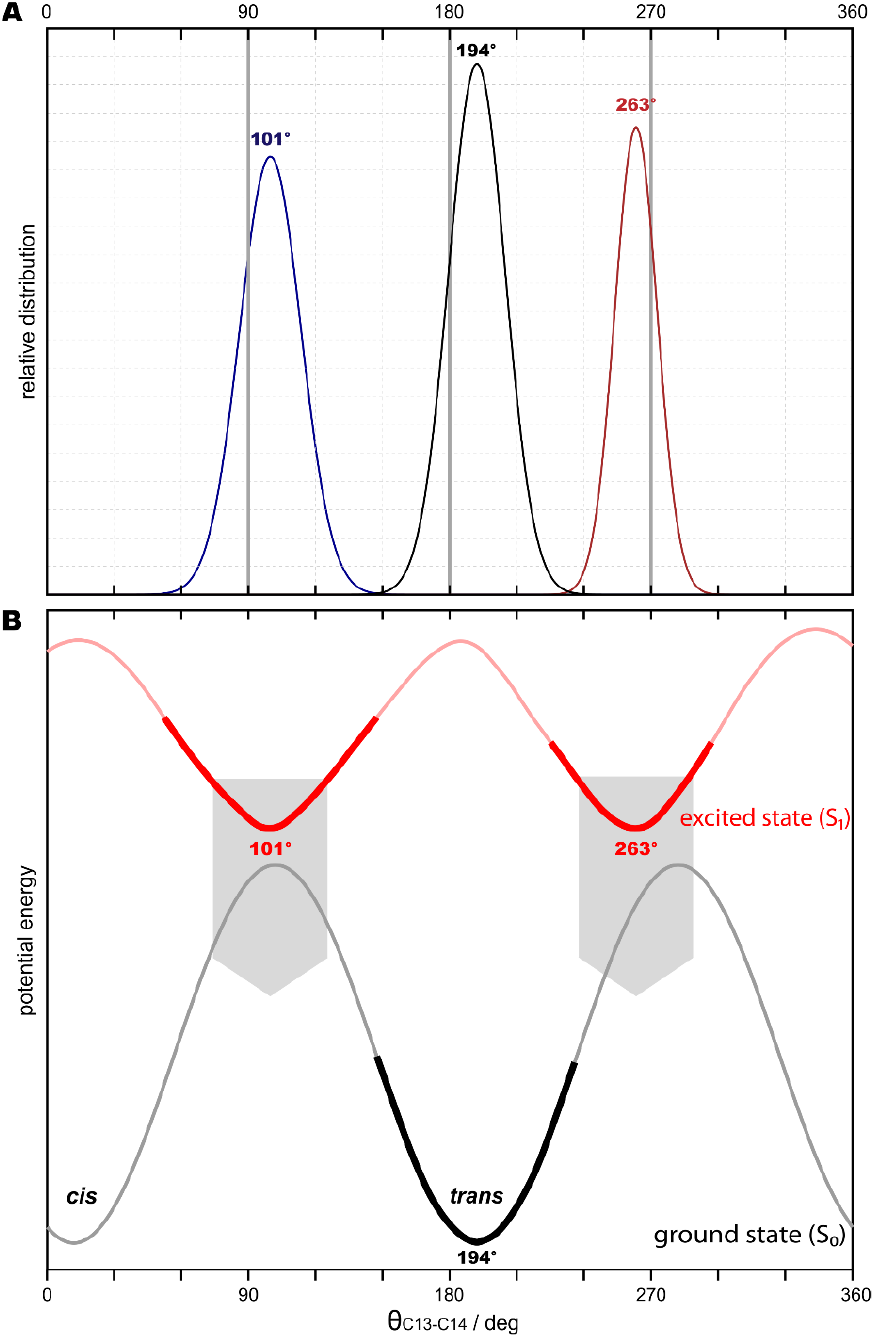
Configurational distribution and anticipated potential energy landscape of retinal in the ground state (S0) and in the excited state (S1). **A** The relative of θ_C13-C14_ angle distribution is given for the equilibrated ground state, step 1 (dotted black curve) and for the state after the 250 ms MD simulation of step 2 with rotation towards either 90° (dotted blue curve) or 270° (dotted red curve), respectively, using a 200 kJ/mol force constant. Gauss-fits of the data reveal a maximum at 194° of the ground state (continuous black line) and 101° (continuous blue line) and 263° (continuous red line) for the excited state, all slightly shifted from the ideal minima. **B** The maxima from A imply minima of the potential energy landscape of the ground state (bold black) and the excited state (bold red). The position of the ground state (gray) and excited state (light red) maxima are hypothetical assumptions. The transitions (gray arrows) from the excited state to the ground state clearly reveal a bias of the achieved retinal configurations depending on the starting position: The 270° rotation equilibrating to 263° leads predominantly to the formation of the *all-trans* ground state, while the 90° rotation equilibrating to 101° leads to the formation of similar amounts of *all-trans* and *cis* configurations (see also Table 1).

Based on the excited state potential minima shifts compared to the ground state maxima an asymmetric distribution of the resulting isomerization states depending on the rotation direction of θ_C13-C14_ towards 90° or 270° would be expected. Indeed, such dependency is observed for the resulting retinal configurations after step 3 summarized in Table 1. The *trans* and *cis* ratio for an excited state restraint θ_C13-C14_ to 90° with 60% to 40% is roughly comparable as visualized (Figure 5B) by the almost aligned excited state potential minimum and ground state maximum. The distribution for the restraint set to 270° is 95% *cis* to 5% *anti* reveals a clear preference for *trans* state. This preference fully agrees with the obviously shifted excited state potential minima towards the *trans* ground state minimum.

In the following step 3, ground state parameters were applied, and 250 fs MM simulations were performed for relaxation. Figure 6 illustrates the resulting overall distribution of θ_C13-C14_ of the in total 2640 isomerization calculations detailed in Table S7. Table 1 summarizes the percentage distribution of the retinal configurations. While the experimentally measured photochemical quantum yield is approximately 60 %, our calculations result in 22% *cis* [45].

**Figure 6:**
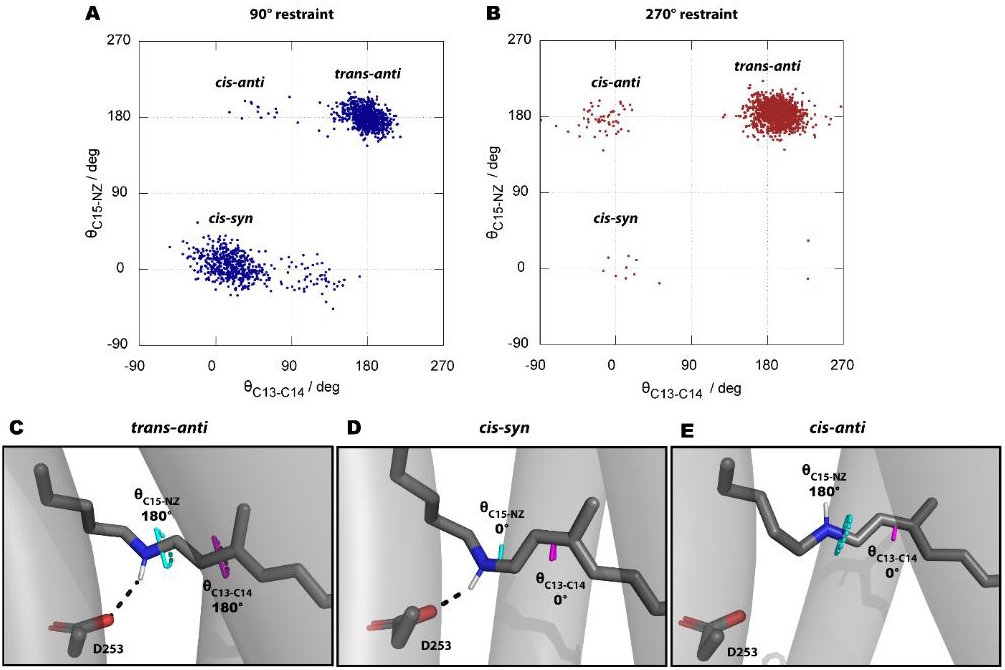
Retinal configurational distribution after isomerization calculation. Distribution plots of θ_C13-C14_ (magenta) and θ_NZ-C15_ (cyan) after relaxation to the ground state (step 3) from the excitations based on an excited state restraint θ_C13-C14_ of 90° (**A**) or 270° (**B**). All values of the distribution plot are given in Table S5. **C** Representative *Cr*ChR2 geometry with all-*trans* retinal configuration. **D** Representative *Cr*ChR2 geometry with 13-*cis* CN-*syn* RSBH^+^ isomerization configuration. **E** Representative *Cr*ChR2 geometry of the 13-*cis* CN-*anti* RSBH^+^ isomerization configuration.

We assume that this deviation would become smaller with more sampling. The need for higher sampling rates to obtain better statistics becomes also evident from the observed strong dependence of the isomerization result on the highly dynamic ground state conformational condition of the protein. For example, the results from the 90° restraint initiated from the geometry at 1,000 ns of monomer A from run 3 isomerizes in 95 % to *cis*, whereas the calculations initiated with the geometry after 950 ns did not show any isomerization at all (see Figure 4). Over-interpretation of the calculated statistics of isomerization configurations should therefore be avoided as the numbers are somewhat affected by limitations of sampling of initial geometries. In addition to the usual sampling limitation, there is a strong dependence on the initial geometries. However, the general conclusion is justified that those distinct retinal configurations computationally observed after photoisomerization qualitatively exist in nature and their distributions are predictable after photochemical transition.

Besides the *trans-anti* configuration, the second most observed configuration is *cis-syn* (20 %). Notably *cis-syn* is almost only observed with an excited state restraint θ_C13-C14_ of 90° (39 %), while with an excited state restraint θ_C13-C14_ of 270° we observed only 1 %. The *cis-anti* configuration is overall observed with 2 % with a predominant fraction for the excited state restraint θ_C13-C14_ of 270° (4 %) and 1 % observed with an excited state restraint θ_C13-C14_ of 90°. These results are in line with experimental IR spectroscopy results [13] indicating a split *Cr*ChR2 photo cycle containing both retinal configurations, *cis-anti* associated with the P_500_ (K) intermediate and *cis-syn* associated with the P_480_ intermediate. [57] This agreement of experiment and computation shows the successful application to obtain spontaneously different isomerization states without biasing a specific configuration.

### 2.5 Ground state force constant dependency control

Due to the outlined natural dependence of MM simulations on the force field parameters we performed a dependency control of our results on the retinal parameter set. There are two different ways to derive retinal parameters for the OPLS/AA force field which are equally valid. One is the method described above following the procedure of Weiner et al.[60] and the other is to derive the force constant for the torsions based on quantum chemical calculations as described by Jorgensen et al.[61].To check if our here presented results depend on the parameterization strategy, we also derived retinal parameters employing the second strategy. Therefore, we compared the torsional barriers of θ_C13-C14_ and θ_C15-NZ_ obtained from quantum mechanical calculations (PBE/6-31G*) with the RSBH^+^ force field parameters, derived in this work. The QM method calculated a torsional barrier of 74.95 kJ/mol for the θ_C15-NZ_ and 93.38 kJ/mol for θ_C13-C14_. The interpolated force constants lead to amplitudes for the MM simulation with 80 kJ/mol for θ_C15-NZ_ and 44.77 kJ/mol for θ_C13-C14_. The MM force field’s value for the θ_C15-NZ_ closely matched the QM result, indicating good agreement in this case. However, the θ_C13-C14_ showed a substantial deviation, with the force constant being approximately half of the QM value. Notably, the force constants were interpolated via the scheme described in the original amber paper [60], making any adjustments to align the value more closely with the QM results impossible within the context of this procedure. If the force constant for θ_C13-C14_ had been doubled to match the QM value more closely, the force at a 14° torsion angle would have increased by 5.24 kJ/mol. That may have shifted the equilibrium angle slightly towards 180° and thus influenced the balance of *cis*-*syn*/*cis*-*anti* described above. We performed a 350 ns MM simulation of a *Cr*ChR2-dimer as described in the Methods section, but with doubled dihedral force constants to match the QM-values. The results are shown in Figure S7 and indicate that the position of the torsion-minimum is not affected. As expected, the width of the dihedral distribution is decreased for the MM simulations with the stronger force constant. So, all conclusions made to derive the potential energy surface are independent on the choice of the force constant.

## 3. Conclusion

In this work, we developed a novel method to calculate RSBH^+^ isomerization based on MM simulations at the natural time scale of femtoseconds. To improve the quality of the MM simulations we optimized the force field parameters of retinal covalently bound to the opsin via a Schiff base for the OPLS/AA force field by means of quantum chemical calculations. Our parameter set considers the quantum chemically observed bond length alteration pattern and the correct hybridization of all atoms.

We employed our method to study RSBH^+^ isomerization of Channelrhodopsin-2 from *Chlamydomonas reinhardtii* (*Cr*ChR2), one of the most frequently used optogenetic tools. After excitation and relaxation from the excited state, we observed the unbiased production of both RSBH^+^ isomerization states, 13-*cis/* CN-*anti* and 13-*cis*/CN-*syn*, besides the reformation of the non-isomerized *all*-*trans*/CN-*anti* state. While other channelrhodopsins exhibit only a single isomerization product leading to a single photocycle, for *Cr*ChR2 such a branched photocycle was experimentally proven by IR spectroscopy. Beyond this, our computed governs an uneven distribution of the resulting isomerization products. Furthermore, we obtained computational structural models of the photointermediates of the split *Cr*ChR2 photocycle, previously identified experimentally, namely the early P_480_ (*cis*-*syn*) and P_500_ (K, *cis*-*anti*) intermediates. Our models provide the structural basis for future investigations of subsequent photocyclic steps leading to the elucidation of their role in ion conductivity e.g. by means of molecular dynamics simulations.

Based on our isomerization calculations using MM simulations with quantum chemical optimized force field parameters, we have gained a deeper understanding of the photochemical dynamics of the protein cofactor retinal. We anticipate that incorporation of this knowledge into structure refinement approaches will improve the understanding of experimental data. This improvement will also resolve significant discrepancies of protein bound retinal topologies reported to the protein data bank, which are not in accordance with chemical and quantum chemical knowledge.

Moreover, our novel simulation approach, which avoids enforced conformational restraints, presents a more natural representation of the RSBH^+^ isomerization process. Our described methodology bridges the gap between biologically relevant time scales and computational accessibility and provides a robust framework to generally explore molecular mechanisms also of other photo-switchable retinal proteins (i.e. type 1 opsins, or type 2 opsins that act as G-protein coupled receptors) with high spatial and temporal resolution. Application of this method to other retinal proteins will provide insights into the structure function relationships and into protein environmental effects that modulate the retinal behavior. Such detailed mechanistic insights have great potential to improve the design of optogenetic tools for precise for biological manipulation of cellular function in the future.

## 4. Methods

### 4.1 Simulation System Preparation

We performed four 1 µs MM simulations of a *Cr*ChR2-dimer inserted in a 1-palmitoyl-2-oleoyl-snglycero-3-phosphocholine (POPC) membrane under physiological conditions. The simulation system was prepared starting from the *Cr*ChR2 X-ray structure with the PDB-ID 6EID [62] using the software package MAXIMOBY (CHEOPS, Germany). The unit cell of the crystal contains two antiparallel *Cr*ChR2 monomers. Using PyMOL, neighboring symmetric units were calculated to generate the *Cr*ChR2 dimer. MAXIMOBY automatically detects potential disulfide bridges and generates interdimer bonds between the cysteines C34 and C36 of each monomer. Afterwards missing side-chain atoms (in chain A in residues Q33, Y35, W75, K76, T78, K103, R147, K205, R207, E273; in chain B in residues D32, K76, T78, K103, R147, R207, D280, I281) were added. The structure was protonated based on the local pKa values of each residue calculated at a pH value of 7 using standard pks-values following Nielsen and Vriend [63]. The amino acid E90 and D156 were protonated as observed by FTIR spectroscopic measurements [13, 50]. We used the Gromacs program suite version 2021 [64] in combination with lambada [65] to place the protein into a POPC bilayer. Water molecules of the first and second solvation shell of the protein membrane complex were set using the solvation approached implemented in MAXIMOBY that is based on the principles of the Vedani algorithm ^64^. To prevent self-interactions of the protein due to periodic boundary conditions within the MM simulation, a cubic simulation box with the dimensions 125*125*125 nm was set and filled with water molecules, sodium and calcium ions, at physiological conditions using the solvation workflow implemented in GROMACS 2021 [64]. Bulk water is provided in cubes and steric clashes may occur at transitions between incomplete cubes or at transitions between the second solvation shell and the bulk-cubes. The clashes were locally resolved through energy optimization in MAXIMOBY using the TIP3P water model [66] and the original Amber84 united atom force field [60]. For the simulation runs with the OPLS/AA force field, the TIP3P water model is converted into the TIP4P water model.

### 4.2 MM Simulation Details

First, the system was heated to room temperature (300 K) in 1 ns with a step size of 1 fs within a nVT simulation, meaning the number of atoms (n), the volume (V), and the temperature (T) is constant while the pressure (p) is flexible. The temperature was kept constant using a V-rescale thermostat [67] with a coupling constant 0.1 ps. The heating was performed in two steps, over the first 100 ps the temperature was raised continuously from 0 K to 100 K, in the following 900 ps the temperature was raised continuously to 300 K. The heating procedure was followed by a 1 ns nVT simulation with a step size of 1 fs under the same conditions as for the heating but at 300 K. Next, we performed a 10 ns npT run with a step size of 1 fs in which the number of atoms, pressure and temperature remained constant using a Berendsen barostat [68] (coupling constant 0.5 ps) and a V-rescale thermostat (coupling constant 0.1 ps) while the volume was kept flexible. Following these equilibration steps we performed three 1 µs npT production runs (step size 2 fs). In the production run, the temperature was controlled by a Nosé-Hoover thermostat [69, 70] (coupling constant 0.5 ps) and a Parrinello-Rahman barostat [71] (coupling constant 2.5 ps) which perform better for equilibrated systems. The equilibrated second half of the trajectory namely the last 500 ns were used for further analysis. The decision for the equilibrated part was made based on the root mean square deviation (RMSD) (see Figure S4) and contact analysis.

### 4.3 Simulation Evaluation

To assess the structural stability and conformational dynamics of the system during the MM simulations, we calculated the root mean square deviation (RMSD) of the atomic positions relative to the initial structure before heating. RMSD is a standard measure for quantifying the average displacement of atoms between the simulated structures and a reference conformation.

For the RMSD calculations, we used the alignment function of MAXIMOBY. Prior to the RMSD analysis, the system’s trajectory was superimposed to the initial structure to remove any translational or rotational motion, ensuring that only internal structural changes were measured. The RMSD was computed for Cα-atoms to track the overall protein stability while minimizing noise from flexible side chains. The RMSD at time *t* was calculated according to the following equation 1:

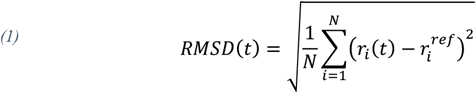

where *N* is the number of atoms selected for analysis, *r*_*i*_ (*t*) is the position of atom *i* at time *t*, and 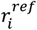 is the position of atom *i* in the reference conformation (here the initial conformation).

The trajectory was sampled at intervals of 0.1 ns and RMSD values were plotted as a function of simulation time. To identify any significant conformational changes or stabilization of the structure, we visually inspected the RMSD plot. A plateau in the RMSD values was taken as an indicator of structural stability.

To monitor more detailed conformational changes and changes in the interaction network we performed a contact analysis implemented in MAXIMOBY. It distinguishes between sidechain and backbone contacts, as well as between hydrogen bond/ionic contacts, simple van der Waals contacts, and van der Waals contacts involving aromatic. The implemented contact analysis algorithm further performs chirality and secondary structure conformation analyses using a combination of Define Secondary Structure of Proteins (DSSP)-analysis [72] and dihedral angle measurements. For all four MM simulations, contact analysis was performed over the complete simulation length of 1 µs, with a resolution of 0.1 ns.

### 4.5 Retinal Force Field Parametrization

For the system preparation we use the original Amber84 united atom force field [60] and for the subsequent molecular mechanics simulations the OPLS/AA force field [61]. The OPLS force field is based on the AMBER force fields and therefore both share the same bonded parameters (bond lengths, bond angles, and torsions), which are derived from experimental and quantum mechanical data. However, they differ significantly in their treatment of non-bonded interactions (electrostatics and van der Waals forces). OPLS fine-tunes its non-bonded parameters using experimental data from liquid simulations to accurately reproduce properties like densities and heats of vaporization. This approach optimizes OPLS for simulations involving organic liquids and small molecules [66, 73]. As we are studying a transmembrane helix protein with focus on the covalently bound small molecule retinal we employed the OPLS/AA force field. Usually only the standard amino acids are provided by default with the force field. For all hetero groups and non-standard residues, we need to add parameters derived by others if available or derive them by ourselves. As to our knowledge no published parameters for the OPLS/AA force field exist we derived a new parameter set as detailed in the Supporting information.

For the POPC lipids we used the parameters from Berger et al. [74] for the OPLS/AA force field. To set up the simulation system, especially with respect to a proper solvation of the membrane, we used OPLS/AA POPC parameters simply by changing the center types from the OPLS/AA to the Amber UA force field. We did not adapt the charges since this was not necessary as we did not perform MM simulations or energy optimizations of the POPC membrane in Amber UA. The center types assured that all POPC lipids were suitably solvated.

## 5. Acknowledgements

This work was supported by Deutsche Forschungsgemeinschaft (DFG, German Research Foundation) Individual Research Grant, “Molecular mechanisms of cation and anion-conducting channelrhodopsins” (GE 599/23-1) to K.G. and the DFG Priority Program SPP1926 (GE 599/19-2 and GE 599/19-1) to K.G.

## 6. Author contributions

**Philipp Althoff**: Formal analysis, Investigation, Methodology, Software, Validation, Visualization, Writing – original draft, Writing – review & editing. **Kristin Labudda**: Formal analysis, Validation, Visualization, Writing – original draft, Writing – review & editing. **Udo Höweler**: Conceptualization, Methodology, Software, Supervision, Writing – original draft, Writing – review & editing. **Mathias Lübben**: Conceptualization, Supervision, Validation, Visualization, Writing – original draft, Writing – review & editing. **Klaus Gerwert**: Funding acquisition, Resources, Supervision, Writing – review & editing. **Carsten Kötting**: Conceptualization, Funding acquisition, Methodology, Resources, Supervision, Validation, Visualization, Writing – original draft, Writing – review & editing. **Till Rudack**: Conceptualization, Data curation, Funding acquisition, Methodology, Project administration, Resources, Software, Supervision, Writing – original draft, Writing – review & editing

## References

1. Briggs, W.R., Spudich, J.L.: Handbook of Photosensory Receptors. Wiley (2005)

2. Ernst, O.P., Lodowski, D.T., Elstner, M., Hegemann, P., Brown, L.S., Kandori, H.: Microbial and animal rhodopsins: structures, functions, and molecular mechanisms. Chemical reviews 114(1), 126–163 (2014). doi: 10.1021/cr4003769

3. Govorunova, E.G., Sineshchekov, O.A., Li, H., Spudich, J.L.: Microbial Rhodopsins: Diversity, Mechanisms, and Optogenetic Applications. Annual review of biochemistry 86, 845–872 (2017). doi: 10.1146/annurev-biochem-101910-144233

4. Govorunova, E.G., Sineshchekov, O.A., Rodarte, E.M., Janz, R., Morelle, O., Melkonian, M., Wong, G.K.-S., Spudich, J.L.: The Expanding Family of Natural Anion Channelrhodopsins Reveals Large Variations in Kinetics, Conductance, and Spectral Sensitivity. Scientific reports 7(1), 43358 (2017). doi: 10.1038/srep43358

5. Zemelman, B.V., Lee, G.A., Ng, M., Miesenböck, G.: Selective photostimulation of genetically chARGed neurons. Neuron 33(1), 15–22 (2002). doi: 10.1016/S0896-6273(01)00574-8

6. Deisseroth, K.: Optogenetics. Nature methods 8(1), 26–29 (2011). doi: 10.1038/nmeth.f.324

7. Wietek, J., Broser, M., Krause, B.S., Hegemann, P.: Identification of a Natural Green Light Absorbing Chloride Conducting Channelrhodopsin from Proteomonas sulcata. The Journal of biological chemistry 291(8), 4121–4127 (2016). doi: 10.1074/jbc.M115.699637

8. Kim, C.K., Adhikari, A., Deisseroth, K.: Integration of optogenetics with complementary methodologies in systems neuroscience. Nature Reviews Neuroscience 18(4), 222–235 (2017). doi: 10.1038/nrn.2017.15

9. Boyden, E.S., Zhang, F., Bamberg, E., Nagel, G., Deisseroth, K.: Millisecond-timescale, genetically targeted optical control of neural activity. Nature neuroscience 8(9), 1263–1268 (2005). doi: 10.1038/nn1525

10. Govorunova, E.G., Sineshchekov, O.A., Spudich, J.L.: Three Families of Channelrhodopsins and Their Use in Optogenetics (review). Neuroscience and Behavioral Physiology 49(2), 163–168 (2019). doi: 10.1007/s11055-019-00710-6

11. Kandori, H.: Biophysics of rhodopsins and optogenetics. Biophysical reviews 12(2), 355– 361 (2020). doi: 10.1007/s12551-020-00645-0

12. Nagel, G., Ollig, D., Fuhrmann, M., Kateriya, S., Musti, A.M., Bamberg, E., Hegemann, P.: Channelrhodopsin-1: a light-gated proton channel in green algae. Science (New York, N.Y.) 296(5577), 2395–2398 (2002). doi: 10.1126/science.1072068

13. Kuhne, J., Vierock, J., Tennigkeit, S.A., Dreier, M.-A., Wietek, J., Petersen, D., Gavriljuk, K., El-Mashtoly, S.F., Hegemann, P., Gerwert, K.: Unifying photocycle model for light adaptation and temporal evolution of cation conductance in channelrhodopsin-2. Proceedings of the National Academy of Sciences of the United States of America 116(19), 9380–9389 (2019). doi: 10.1073/pnas.1818707116

14. Zhang, Y.-P., Oertner, T.G.: Optical induction of synaptic plasticity using a light-sensitive channel. Nature methods 4(2), 139–141 (2007). doi: 10.1038/nmeth988

15. Petreanu, L., Huber, D., Sobczyk, A., Svoboda, K.: Channelrhodopsin-2-assisted circuit mapping of long-range callosal projections. Nature neuroscience 10(5), 663–668 (2007). doi: 10.1038/nn1891

16. Schneider, F., Grimm, C., Hegemann, P.: Biophysics of Channelrhodopsin. Annual review of biophysics 44, 167–186 (2015). doi: 10.1146/annurev-biophys-060414-034014

17. Arenkiel, B.R., Peca, J., Davison, I.G., Feliciano, C., Deisseroth, K., Augustine, G.J., Ehlers, M.D., Feng, G.: In vivo light-induced activation of neural circuitry in transgenic mice expressing channelrhodopsin-2. Neuron 54(2), 205–218 (2007). doi: 10.1016/j.neuron.2007.03.005

18. Douglass, A.D., Kraves, S., Deisseroth, K., Schier, A.F., Engert, F.: Escape behavior elicited by single, channelrhodopsin-2-evoked spikes in zebrafish somatosensory neurons. Current biology : CB 18(15), 1133–1137 (2008). doi: 10.1016/j.cub.2008.06.077

19. Gradinaru, V., Mogri, M., Thompson, K.R., Henderson, J.M., Deisseroth, K.: Optical deconstruction of parkinsonian neural circuitry. Science (New York, N.Y.) 324(5925), 354– 359 (2009). doi: 10.1126/science.1167093

20. Sahel, J.-A., Boulanger-Scemama, E., Pagot, C., Arleo, A., Galluppi, F., Martel, J.N., Degli Esposti, S., Delaux, A., Saint Aubert, J.-B. de Montleau,, C. de, Gutman, E., Audo, I., Duebel, J., Picaud, S., Dalkara, D., Blouin, L., Taiel, M., Roska, B.: Partial recovery of visual function in a blind patient after optogenetic therapy. Nature medicine 27(7), 1223– 1229 (2021). doi: 10.1038/s41591-021-01351-4

21. Shimano, T., Fyk-Kolodziej, B., Mirza, N., Asako, M., Tomoda, K., Bledsoe, S., Pan, Z.H., Molitor, S., Holt, A.G.: Assessment of the AAV-mediated expression of channelrhodopsin-2 and halorhodopsin in brainstem neurons mediating auditory signaling. Brain research 1511, 138–152 (2013). doi: 10.1016/j.brainres.2012.10.030

22. Garczarek, F., Gerwert, K.: Functional waters in intraprotein proton transfer monitored by FTIR difference spectroscopy. Nature 439(7072), 109–112 (2006). doi: 10.1038/nature04231

23. Garczarek, F., Brown, L.S., Lanyi, J.K., Gerwert, K.: Proton binding within a membrane protein by a protonated water cluster. Proceedings of the National Academy of Sciences of the United States of America 102(10), 3633–3638 (2005). doi: 10.1073/pnas.0500421102

24. Gerwert, K., Freier, E., Wolf, S.: The role of protein-bound water molecules in microbial rhodopsins. Biochimica et biophysica acta 1837(5), 606–613 (2014). doi: 10.1016/j.bbabio.2013.09.006

25. Tenboer, J., Basu, S., Zatsepin, N., Pande, K., Milathianaki, D., Frank, M., Hunter, M., Boutet, S., Williams, G.J., Koglin, J.E., Oberthuer, D., Heymann, M., Kupitz, C., Conrad, C., Coe, J., Roy-Chowdhury, S., Weierstall, U., James, D., Wang, D., Grant, T., Barty, A., Yefanov, O., Scales, J., Gati, C., Seuring, C., Srajer, V., Henning, R., Schwander, P., Fromme, R., Ourmazd, A., Moffat, K., van Thor, J.J., Spence, J.C.H., Fromme, P., Chapman, H.N., Schmidt, M.: Time-resolved serial crystallography captures high-resolution intermediates of photoactive yellow protein. Science (New York, N.Y.) 346(6214), 1242–1246 (2014). doi: 10.1126/science.1259357

26. Barends, T.R.M., Foucar, L., Ardevol, A., Nass, K., Aquila, A., Botha, S., Doak, R.B., Falahati, K., Hartmann, E., Hilpert, M., Heinz, M., Hoffmann, M.C., Köfinger, J., Koglin, J.E., Kovacsova, G., Liang, M., Milathianaki, D., Lemke, H.T., Reinstein, J., Roome, C.M., Shoeman, R.L., Williams, G.J., Burghardt, I., Hummer, G., Boutet, S., Schlichting, I.: Direct observation of ultrafast collective motions in CO myoglobin upon ligand dissociation. Science (New York, N.Y.) 350(6259), 445–450 (2015). doi: 10.1126/science.aac5492

27. Pande, K., Hutchison, C.D.M., Groenhof, G., Aquila, A., Robinson, J.S., Tenboer, J., Basu, S., Boutet, S., DePonte, D.P., Liang, M., White, T.A., Zatsepin, N.A., Yefanov, O., Morozov, D., Oberthuer, D., Gati, C., Subramanian, G., James, D., Zhao, Y., Koralek, J., Brayshaw, J., Kupitz, C., Conrad, C., Roy-Chowdhury, S., Coe, J.D., Metz, M., Xavier, P.L., Grant, T.D., Koglin, J.E., Ketawala, G., Fromme, R., Šrajer, V., Henning, R., Spence, J.C.H., Ourmazd, A., Schwander, P., Weierstall, U., Frank, M., Fromme, P., Barty, A., Chapman, H.N., Moffat, K., van Thor, J.J., Schmidt, M.: Femtosecond structural dynamics drives the trans/cis isomerization in photoactive yellow protein. Science (New York, N.Y.) 352(6286), 725–729 (2016). doi: 10.1126/science.aad5081

28. Nango, E., Royant, A., Kubo, M., Nakane, T., Wickstrand, C., Kimura, T., Tanaka, T., Tono, K., Song, C., Tanaka, R., Arima, T., Yamashita, A., Kobayashi, J., Hosaka, T., Mizohata, E., Nogly, P., Sugahara, M., Nam, D., Nomura, T., Shimamura, T., Im, D., Fujiwara, T., Yamanaka, Y., Jeon, B., Nishizawa, T., Oda, K., Fukuda, M., Andersson, R., BÅth, P., Dods, R., Davidsson, J., Matsuoka, S., Kawatake, S., Murata, M., Nureki, O., Owada, S., Kameshima, T., Hatsui, T., Joti, Y., Schertler, G., Yabashi, M., Bondar, A.-N., Standfuss, J., Neutze, R., Iwata, S.: A three-dimensional movie of structural changes in bacteriorhodopsin. Science (New York, N.Y.) 354(6319), 1552–1557 (2016). doi: 10.1126/science.aah3497

29. Nogly, P., Weinert, T., James, D., Carbajo, S., Ozerov, D., Furrer, A., Gashi, D., Borin, V., Skopintsev, P., Jaeger, K., Nass, K., BÅth, P., Bosman, R., Koglin, J., Seaberg, M., Lane, T., Kekilli, D., Brünle, S., Tanaka, T., Wu, W., Milne, C., White, T., Barty, A., Weierstall, U., Panneels, V., Nango, E., Iwata, S., Hunter, M., Schapiro, I., Schertler, G., Neutze, R., Standfuss, J.: Retinal isomerization in bacteriorhodopsin captured by a femtosecond x-ray laser. Science (New York, N.Y.) 361(6398) (2018). doi: 10.1126/science.aat0094

30. Weinert, T., Skopintsev, P., James, D., Dworkowski, F., Panepucci, E., Kekilli, D., Furrer, A., Brünle, S., Mous, S., Ozerov, D., Nogly, P., Wang, M., Standfuss, J.: Proton uptake mechanism in bacteriorhodopsin captured by serial synchrotron crystallography. Science (New York, N.Y.) 365(6448), 61–65 (2019). doi: 10.1126/science.aaw8634

31. Abramson, J., Adler, J., Dunger, J., Evans, R., Green, T., Pritzel, A., Ronneberger, O., Willmore, L., Ballard, A.J., Bambrick, J., Bodenstein, S.W., Evans, D.A., Hung, C.-C., O’Neill, M., Reiman, D., Tunyasuvunakool, K., Wu, Z., Žemgulytė, A., Arvaniti, E., Beattie, C., Bertolli, O., Bridgland, A., Cherepanov, A., Congreve, M., Cowen-Rivers, A.I., Cowie, A., Figurnov, M., Fuchs, F.B., Gladman, H., Jain, R., Khan, Y.A., Low, C.M.R., Perlin, K., Potapenko, A., Savy, P., Singh, S., Stecula, A., Thillaisundaram, A., Tong, C., Yakneen, S., Zhong, E.D., Zielinski, M., Žídek, A., Bapst, V., Kohli, P., Jaderberg, M., Hassabis, D., Jumper, J.M.: Accurate structure prediction of biomolecular interactions with AlphaFold 3. Nature 630(8016), 493–500 (2024). doi: 10.1038/s41586-024-07487-w

32. Baek, M., McHugh, R., Anishchenko, I., Jiang, H., Baker, D., DiMaio, F.: Accurate prediction of protein–nucleic acid complexes using RoseTTAFoldNA. Nature methods 21(1), 117–121 (2024). doi: 10.1038/s41592-023-02086-5

33. Jumper, J., Evans, R., Pritzel, A., Green, T., Figurnov, M., Ronneberger, O., Tunyasuvunakool, K., Bates, R., Žídek, A., Potapenko, A., Bridgland, A., Meyer, C., Kohl, S.A.A., Ballard, A.J., Cowie, A., Romera-Paredes, B., Nikolov, S., Jain, R., Adler, J., Back, T., Petersen, S., Reiman, D., Clancy, E., Zielinski, M., Steinegger, M., Pacholska, M., Berghammer, T., Bodenstein, S., Silver, D., Vinyals, O., Senior, A.W., Kavukcuoglu, K., Kohli, P., Hassabis, D.: Highly accurate protein structure prediction with AlphaFold. Nature 596(7873), 583–589 (2021). doi: 10.1038/s41586-021-03819-2

34. Sen, S., Kar, R.K., Borin, V.A., Schapiro, I.: Insight into the isomerization mechanism of retinal proteins from hybrid quantum mechanics/molecular mechanics simulations. WIREs Comput Mol Sci 12(1) (2022). doi: 10.1002/wcms.1562

35. Tajkhorshid, E., Baudry, J., Schulten, K., Suhai, S.: Molecular Dynamics Study of the Nature and Origin of Retinal’s Twisted Structure in Bacteriorhodopsin. Biophysical journal 78(2), 683–693 (2000). doi: 10.1016/S0006-3495(00)76626-4

36. Perilla, J.R., Goh, B.C., Cassidy, C.K., Liu, B., Bernardi, R.C., Rudack, T., Yu, H., Wu, Z., Schulten, K.: Molecular dynamics simulations of large macromolecular complexes. Current opinion in structural biology 31, 64–74 (2015). doi: 10.1016/j.sbi.2015.03.007

37. Bogomolni, R.A., Spudich, J.L.: Identification of a third rhodopsin-like pigment in phototactic Halobacterium halobium. Proceedings of the National Academy of Sciences of the United States of America 79(20), 6250–6254 (1982). doi: 10.1073/pnas.79.20.6250

38. Takahashi, T., Mochizuki, Y., Kamo, N., Kobatake, Y.: Evidence that the long-lifetime photointermediate of s-rhodopsin is a receptor for negative phototaxis in Halobacterium halobium. Biochemical and biophysical research communications 127(1), 99–105 (1985). doi: 10.1016/S0006-291X(85)80131-5

39. Sineshchekov, O.A., Jung, K.-H., Spudich, J.L.: Two rhodopsins mediate phototaxis to low- and high-intensity light in Chlamydomonas reinhardtii. Proceedings of the National Academy of Sciences of the United States of America 99(13), 8689–8694 (2002). doi: 10.1073/pnas.122243399

40. Sineshchekov, O.A., Govorunova, E.G., Jung, K.-H., Zauner, S., Maier, U.-G., Spudich, J.L.: Rhodopsin-mediated photoreception in cryptophyte flagellates. Biophysical journal 89(6), 4310–4319 (2005). doi: 10.1529/biophysj.105.070920

41. Mongodin, E.F., Nelson, K.E., Daugherty, S., Deboy, R.T., Wister, J., Khouri, H., Weidman, J., Walsh, D.A., Papke, R.T., Sanchez Perez, G., Sharma, A.K., Nesbø, C.L., MacLeod, D., Bapteste, E., Doolittle, W.F., Charlebois, R.L., Legault, B., Rodriguez-Valera, F.: The genome of Salinibacter ruber: convergence and gene exchange among hyperhalophilic bacteria and archaea. Proceedings of the National Academy of Sciences of the United States of America 102(50), 18147–18152 (2005). doi: 10.1073/pnas.0509073102

42. Govorunova, E.G., Sineshchekov, O.A., Spudich, J.L.: Emerging Diversity of Channelrhodopsins and Their Structure-Function Relationships. Frontiers in cellular neuroscience 15, 800313 (2021). doi: 10.3389/fncel.2021.800313

43. Spudich, J.L., Sineshchekov, O.A., Govorunova, E.G.: Mechanism divergence in microbial rhodopsins. Biochimica et biophysica acta 1837(5), 546–552 (2014). doi: 10.1016/j.bbabio.2013.06.006

44. R. W. Schoenlein, L. A. Peteanu, Q. Wang, R. A. Mathies, C. V. Shank: Femtosecond dynamics of cis-trans isomerization in a visual pigment analog: isorhodopsin. J. Phys. Chem. (The Journal of Physical Chemistry)(97), 12087–12092 (1993)

45. Klaus Schulten, Shigehiko Hayashi: Quantum Biology of Retinal (2014)

46. K. W. Foster, J. Saranak, N. Patel, G. Zarilli, M. Okabe, T. Kline, and K. Nakanishi.: A rhodopsin is the functional photoreceptor for phototaxis in the unicellular eukaryote Chlamydomonas. Nature (311), 756–759 (1984)

47. Hegemann, P., Gärtner, W., Uhl, R.: All-trans retinal constitutes the functional chromophore in Chlamydomonas rhodopsin. Biophysical journal 60(6), 1477–1489 (1991). doi: 10.1016/S0006-3495(91)82183-X

48. Lawson, M.A., Zacks, D.N., Derguini, F., Nakanishi, K., Spudich, J.L.: Retinal analog restoration of photophobic responses in a blind Chlamydomonas reinhardtii mutant. Evidence for an archaebacterial like chromophore in a eukaryotic rhodopsin. Biophysical journal 60(6), 1490–1498 (1991). doi: 10.1016/S0006-3495(91)82184-1

49. Nagel, G., Szellas, T., Kateriya, S., Adeishvili, N., Hegemann, P., Bamberg, E.: Channelrhodopsins: directly light-gated cation channels. Biochemical Society transactions 33(Pt 4), 863–866 (2005). doi: 10.1042/BST0330863

50. Lórenz-Fonfría, V.A., Resler, T., Krause, N., Nack, M., Gossing, M., Fischer von Mollard, G., Bamann, C., Bamberg, E., Schlesinger, R., Heberle, J.: Transient protonation changes in channelrhodopsin-2 and their relevance to channel gating. Proceedings of the National Academy of Sciences of the United States of America 110(14), E1273–81 (2013). doi: 10.1073/pnas.1219502110

51. Arieh Warshel: Bicyle-pedal model for the first step in the vision process. Nature (260), 679–683 (1976)

52. M. Nonella, A. Windemuth, K. Schulten: STRUCTURE OF BACTERIORHODOPSIN and in situ ISOMERIZATION OF RETINAL: A MOLECULAR DYNAMICS STUDY*. Photochemistry and photobiology (54 (6)), 937–948 (1991)

53. Sen, S., Kar, R.K., Borin, V.A., Schapiro, I.: Insight into the isomerization mechanism of retinal proteins from hybrid quantum mechanics/molecular mechanics simulations. WIREs Comput Mol Sci 12(1) (2022). doi: 10.1002/wcms.1562

54. Ardevol, A., Hummer, G.: Retinal isomerization and water-pore formation in channelrhodopsin-2. Proceedings of the National Academy of Sciences of the United States of America 115(14), 3557–3562 (2018). doi: 10.1073/pnas.1700091115

55. Tribello, G.A., Bonomi, M., Branduardi, D., Camilloni, C., Bussi, G.: PLUMED 2: New feathers for an old bird. Computer Physics Communications 185(2), 604–613 (2014). doi: 10.1016/j.cpc.2013.09.018

56. Xin, Q., Zhang, W., Yuan, S.: The Mechanism of the Channel Opening in Channelrhodopsin-2: A Molecular Dynamics Simulation. International Journal of Molecular Sciences 24(6) (2023). doi: 10.3390/ijms24065667

57. Bellucci, L., Capone, M., Daidone, I., Zanetti-Polzi, L.: Conformational heterogeneity and protonation equilibria shape the photocycle branching in channelrhodopsin-2. International journal of biological macromolecules, 140977 (2025). doi: 10.1016/j.ijbiomac.2025.140977

58. Hermone, A., Kuczera, K.: Free-Energy Simulations of the Retinal Cis → Trans Isomerization in Bacteriorhodopsin. Biochemistry 37(9), 2843–2853 (1998). doi: 10.1021/bi9717789

59. Kunishige, S., Katori, T., Baba, M., Nakajima, M., Endo, Y.: Spectroscopic study on deuterated benzenes. I. Microwave spectra and molecular structure in the ground state. The Journal of Chemical Physics 143(24), 244302 (2015). doi: 10.1063/1.4937949

60. Weiner, S.J., Kollman, P.A., Case, D.A., Singh, U.C., Ghio, C., Alagona, G., Profeta, S., Weiner, P.: A new force field for molecular mechanical simulation of nucleic acids and proteins. J. Am. Chem. Soc. 106(3), 765–784 (1984). doi: 10.1021/ja00315a051

61. Jorgensen, W.L., Maxwell, D.S., Tirado-Rives, J.: Development and Testing of the OPLS All-Atom Force Field on Conformational Energetics and Properties of Organic Liquids. J. Am. Chem. Soc. 118(45), 11225–11236 (1996). doi: 10.1021/ja9621760

62. Volkov, O., Kovalev, K., Polovinkin, V., Borshchevskiy, V., Bamann, C., Astashkin, R., Marin, E., Popov, A., Balandin, T., Willbold, D., Büldt, G., Bamberg, E., Gordeliy, V.: Structural insights into ion conduction by channelrhodopsin 2. Science (New York, N.Y.) 358(6366) (2017). doi: 10.1126/science.aan8862

63. J. E. Nielsen G. V.: Optimizing the hydrogen-bond network in Poisson–Boltzmann equation-based pKa calculations. Proteins (43), 403–412 (2001)

64. Abraham, M.J., Murtola, T., Schulz, R., Páll, S., Smith, J.C., Hess, B., Lindahl, E.: GROMACS: High performance molecular simulations through multi-level parallelism from laptops to supercomputers. SoftwareX 1-2, 19–25 (2015). doi: 10.1016/j.softx.2015.06.001

65. Schmidt, T.H., Kandt, C.: LAMBADA & InflateGRO2: Efficient Membrane Alignment and Insertion of Membrane Proteins for Molecular Dynamics Simulations. Biophysical journal 102(3), 173a (2012). doi: 10.1016/j.bpj.2011.11.938

66. Jorgensen, W.L., Chandrasekhar, J., Madura, J.D., Impey, R.W., Klein, M.L.: Comparison of simple potential functions for simulating liquid water. The Journal of Chemical Physics 79(2), 926–935 (1983). doi: 10.1063/1.445869

67. Bussi, G., Donadio, D., Parrinello, M.: Canonical sampling through velocity rescaling. The Journal of Chemical Physics (126 (1)), 14101 (2007). doi: 10.1063/1.2408420

68. Berendsen, H.J.C., van Postma, J.P., van Gunsteren, W.F., DiNola, A., Haak, J.R.: Molecular dynamics with coupling to an external bath. The Journal of Chemical Physics 81(8), 3684–3690 (1984)

69. Nosé, S.: A unified formulation of the constant temperature molecular dynamics methods. The Journal of Chemical Physics 81(1), 511–519 (1984)

70. Hoover, W.G.: Canonical dynamics: Equilibrium phase-space distributions. Physical review A 31(3), 1695 (1985)

71. Parrinello, M., Rahman, A.: Polymorphic transitions in single crystals: A new molecular dynamics method. Journal of Applied physics 52(12), 7182–7190 (1981)

72. Kabsch, W., Sander, C.: Dictionary of protein secondary structure: pattern recognition of hydrogen-bonded and geometrical features. Biopolymers 22(12), 2577–2637 (1983). doi: 10.1002/bip.360221211

73. Jorgensen, S.: Optimized intermolecular potential functions for amides and peptides. Structure and properties of liquid amides. J. Am. Chem. Soc.(107), 569–578 (1985)

74. Berger, O., Edholm, O., Jähnig, F.: Molecular dynamics simulations of a fluid bilayer of dipalmitoylphosphatidylcholine at full hydration, constant pressure, and constant temperature. Biophysical journal 72(5), 2002–2013 (1997). doi: 10.1016/S0006-3495(97)78845-3

