## Supplemental text, tables and figures for "A fast and accurate calculation method for light induced isomerization of retinal proteins in real time"

**Supporting Information**

#### Retinal Parameterization Strategy

The accuracy of force field parameters is a key for valid molecular mechanics simulations. For all hetero groups and non-standard residues, we need to add parameters to the force field derived by others if available or derive them by ourselves. As to our knowledge no published parameters for the OPLS/AA force field exist we derived a new parameter set as detailed in the following.

Since $RSBH^+$ is covalently bound to the protein, a QC system was introduced for the derivation of parameters. OPLS/AA extends the united atom OPLS force field by the explicit contemplation of hydrogen atoms and the Refinement of torsional parameters by regarding RHF/6-31G* calculations [1]. For Amber the charges are based on ESP-fits with RHF/STO3G [2]. Therefore, we used QC calculations following the strategy described by Weiner et al. [3] for the Amber UA force field and adapted it to the OPLS/AA force field. First QC systems were carefully defined to ensure a smooth transition between the existing lysine-parameters and the newly developed retinal-parameters, which is crucial because retinal is covalently bound to its surrounding environment. Bonded and dihedral interactions were parameterized according to Amber force field standards, ensuring compatibility with existing force field frameworks. The atomic charges were obtained by quantum-mechanical calculations. Crucial for this step was a performed QM charge calculation on an uncharged carbon-based QC system, which allowed a comparison to existing hydrocarbon parameters (see Figure S8). Lennard-Jones parameters (sigma and epsilon) were taken directly from the standard values of the OPLS-AA force field.

#### QC systems

First, we identified a QC system for the quantum chemical calculations on $RSBH^+$ to derive bonded parameters and partial charges. The decision for the quantum chemical system is already a crucial step as $RSBH^+$ is a heterogroup that has a covalent bond to the protein, built by a nucleophile addition of a sidechain-amine of lysine and retinal forming an imine. Since lysine as a standard amino acid is already parametrized, we decided to set the transition between the existing parameters and our newly introduced retinal parameters at the bond between CD and CE. Figure S9 shows that lysine is split into two charge groups within the OPLS/AA force field. CE, NZ and their corresponding hydrogens form a group with a charge of +1e and the rest of the lysine atoms form a second group with a charge of 0e and in the following referred as lysine core group. The charged group will be replaced by our QC system, which also contains a net charge of +1e. Figure S1 shows both groups.

The lysine-core group remains unchanged while the QC system will be used to determine the missing parameters. As the two systems are connected by a covalent bond a hydrogen atom that saturates the CE atom in our QC system is introduced (see Figure S1). The treatment of this saturation-hydrogen and the transition between QC system and lysine-core group will be explained in each part separately. Figure S3A shows the QC system for $RSBH^+$. We also performed calculations on RSB, for the validation of the conjugated π-electron system. The QC system of RSB is also shown in Figure S3B.

**Bonded Parameters**

As the bond conjugation of the retinal is very crucial we first checked the overall geometrical properties of $RSBH^+$. With quantum chemical accuracy energy optimizations of the QC systems at different quantum chemical levels were performed using Gaussian09 [4]. We compared results using the functionals RHF [5], B3LYP [6], PBE [7], and MP2 [8] in combination with the 6-31G* basis-set. The multiplicity was set to 1 while the charge was 1e for $RSBH^+$ and 0e for RSB respectively. Table S2 shows the comparison of the calculated bond-lengths along the conjugated π -system for $RSBH^+$. While all higher level methods (B3LYP, MP2 and PBE) are constituent revealing bond length equalization for the bonds from C15-C14 to C10-C9, the RHF geometry exhibits a clearly pronounced bond length alteration. The RHF results are not further considered since the delocalized π -electron system is not described correctly by this method and the results of the other 3 are visualized in Figure S10 through a $RSBH^+$ with a revised structural formula. We also performed PBE/6-31G* calculations for the RSB. Table S8 compares the PBE-results for $RSBH^+$ and RSB. RSB shows a clearly pronounced bond-length alteration among all bonds in contrast to $RSBH^+$.

The following key features were observed from the quantum chemical calculations of $RSBH^+$. The NZ-C15 bond shows a length of 1.34 Å. The following six C-C bonds exhibited nearly equal bond lengths of 1.4 Å. Bond C8-C9 has a more pronounced single-bond character of 1.43 Å, followed by the bond C7-C8 with a more pronounced double-bond character of 1.38 Å. Bond C6-C7 has the longest length with 1.44 Å and C5-C6 exhibits again 1.38 Å.

The QC results indicate that the conjugated π-electron system extends from NZ-C15 to C10-C9, with bond lengths alternating afterwards. We assign a bond length of 1.4 Å to all carbon-carbon bonds in the conjugated system for simplicity. Table S1 displays the assigned bond length distribution in our OPLS/AA parameters, reflecting specific variations in bond lengths across the conjugated region based on our QC results. For the Transition between QC system and lysine core group, namely the CE-NZ bond, the values from the standard OPLS/AA-Lysine parameterization were used.

As mentioned in the introduction, OPLS/AA is based on the amber force field and bonded interactions are described in the same way. Force constants for new introduced C-C and C-N bonds in OPLS/AA, are therefore obtained by application of the regulation for new parameters in the amber force field as follows: For C-C bonds the force constant must be interpolated between the value for single C-C Bonds of 265265 kJ/mol/nm$^2$ at 1.507 Å and the value for double C-C Bonds of 476976 kJ/mol/nm$^2$ at 1.336 Å. The same regulations apply for CN-bonds witch values of 476976 kJ/mol/nm$^2$ at 1.273 Å and 282001 kJ/mol/nm$^2$ at 1.449 Å. Figure S11 represents this approach graphically, while Table S1 contains the derived bond parameters.

**Bond angles**

All angle potentials in the π-system were derived from Phenylalanine. Therefore, the bond angles are set to 120° and the force constant to 527,184 kJ/mol/$\Phi^2$. We calculated equilibrium-angles by energy-minimizing our QC system with RHF, B3LYP, PBE and MP2 witch 6-31G* each. Table S9 shows the calculated angles and compares them with MM-minimized structures for the forcefield AMBER and OPLS/AA. MM and QM angles are in general in good agreement, with the highest deviation being ~5° at C8-C7-C6. Table S5 presents all values for RET in OPLS/AA and Table S9 presents QM-calculated angles for the $RSBH^+$.

**Dihedrals**

In OPLS, most dihedrals are adopted from the Amber force field. For developing OPLS/AA torsions, RHF/6-31G* calculations were used to generate torsional profiles. However, the original parametrization did not include charged conjugated π-systems. As seen in the bond analysis, RHF/6-31G* struggles with accurately describing conjugated systems like $RSBH^+$. Therefore, we followed the approach from original Amber and interpolated the dihedral force constants similarly to how we interpolated the bond force constants.

For dihedrals around CC bonds witch lengths between 1.336 Å and 1.4 Å, the force constant is interpolated between 125,52 kJ/mol/rad$^2$ (1.336 Å) and 22.3844 kJ/mol/rad$^2$ (1.4 Å) and for dihedrals around CC-bonds with lengths between 1.4 Å and 1.570 Å the values are 22.3844 kJ/mol/rad$^2$ and 0 kJ/mol/rad$^2$. The dihedral force-constant of CN is interpolated between 125.52 kJ/mol/rad$^2$ at a bond length of 1.273 Å, 41.84 kJ/mol/rad$^2$ at 1.335 Å and 0 kJ/mol/rad$^2$ at 1.449 Å. The Interpolation is represented graphically in Figure S12. Table S6 summarizes all dihedral parameters of the $RSBH^+$. The Multiplicity of double- or conjugated bonds is always 2. $\Phi_0$ was chosen to be 180°, since the $RSBH^+$ is in the *all-trans* configuration in the ground state.

**QM-calculation of Torsion-Potential**

To estimate how the interpolated force constants differ from quantum mechanically determined torsional energy profiles, we conducted PBE/6-31G* calculations on the $RSBH^+$ QC system. The torsions around the C15-NZ bond ($\theta_{C15\text{-}NZ}$) and C13-C14 bond ($\theta_{C13\text{-}C14}$) were systematically analysed by freezing the dihedral angle every 10° and optimizing the structure. The resulting energy differences ($\Delta E$) were compared to the maximal torsional energies of our MM torsion potentials.

**Charges**

In OPLS, the atomic partial charges are fitted not only to quantum mechanically derived electrostatic potentials (ESP) but are also adjusted to reproduce experimental data. Specifically, the OPLS force field uses experimental liquid properties such as densities and heats of vaporization as benchmarks. These experimental values help refine the non-bonded parameters, ensuring that the force field can accurately model intermolecular interactions and the physical behavior of liquid systems, aligning both computational and experimental results. In the absence of data for liquid $RSBH^+$, we performed quantum mechanical calculations for the closest approximation. For the calculation of charge distributions during the development of force fields, commonly used approaches included RHF/6-31G* and RHF/STO-3G methods. These methods employed either Mulliken population analysis or electrostatic potential (ESP) fits to derive atomic charges. We performed calculations with both methods on a carbon framework for validation. The carbon framework contained the same atoms as our QC system, but the nitrogen was substituted by a carbon atom, resulting in a total charge of 0e for this system. The uncharged carbon framework is a less complex system and allows comparison to Phenylalanine or Ethylene-based amino acids. The structure was optimized previously with PBE/6-31G*. The results are presented in Table S10.

To determine the best method for approximating OPLS/AA charges, a comparison with standard OPLS/AA charges is essential. Typically, hydrogen atoms in hydrocarbons carry a charge of +0.060e, and carbon atoms in methyl groups carry -0.180e, resulting in a net neutral charge. Substitutions of hydrogens with carbons reduces the charge by 0.060e per bond, making fully carbon-substituted carbons neutral (0e). Table S10 shows that RHF/STO3G Mulliken charges closely match OPLS/AA charges for hydrocarbons, making this method suitable for calculating charges in the absence of experimental data for liquid $RSBH^+$.

Using RHF/STO3G Mulliken charges for $RSBH^+$, we observe polarization in carbons associated with the conjugated $\pi$-system due to the molecule's +1e charge. The nitrogen atom (NZ) has a charge of 0.306e, comparable to protonated lysine (0.300e) in OPLS/AA. Methyl groups and the ring system retain the typical charge of the carbon framework, with +0.060e for hydrogens and -0.180e for carbons. To translate quantum chemistry values to force field values, all charges were globally symmetrized, ensuring consistent charges for methyl/ethyl hydrogens and symmetry-equivalent atoms. Additionally, methyl groups C16 and C17 were set to OPLS/AA-standard values, and the charge distribution was slightly adjusted.

For the CE carbon, the charge of the associated hydrogen was incorporated into the self-charge. Table S3 presents the final retinal (RET) residue charges within the OPLS/AA force field in the last column. For the lysine core group, the OPLS/AA standard values were taken over. Additionally, AMBER UA force field charges are provided, derived from RHF/STO3G ESP fits, where implicit hydrogens have their charges added to the carbons.

#### Lennard Jones

For the Implementation of $RSBH^+$ to the OPLS/AA force field, the Lennard-Jones parameters $\sigma$ (the point at which the potential is zero) and $\varepsilon$ (the depth of the potential well) are derived from standard OPLS/AA values. To carbons of methyl groups and the ring-system the parameter of opls/aa-centertype opls_135 (CT) was assigned with sigma=3.50000e-01 nm and epsilon= 2.76144e-01 kJ/mol, reflecting their alkyl nature. For carbons of the conjugated $\pi$-electron system without connection to a methyl-group the values of opls_145 (CA) with sigma 3.55000e-01 nm and epsilon 3.17984e-01 kJ/mol were assigned and for those with methyl-group the values of parameter opls_143 (CM) with sigma 3.55000e-01 nm and epsilon 2.92889e-01 kJ/mol. All values are presented in Table S3.

**Table S1.** Bond-lengths and force-constant implementation in the OPLS/AA force-field for the conjugated pi-electron system in RSBH+

| **bond** | | | **$r_0$/ Å** | | | **Force constant $k_B$ / (kJ/(mol*nm$^2$))** | | |
|---|---|---|---|---|---|---|---|---|
| CHARMM | Amber | OPLS | CHARMM | Amber | OPLS | CHARMM | Amber | OPLS |
| NZ-C15 | NR-RA | NZ-C15 | 1.296 | 1.33 | 1.340 | 507937.6 | 399320.96 | 402750.8 |
| C15-C14 | RA-RB | C15-C14 | 1.455 | 1.42 | 1.400 | 209200 | 286436.64 | 392459.2 |
| C14-C13 | RB-RC | C14-C13 | 1.327 | 1.39 | 1.400 | 428441.6 | 323088.48 | 392459.2 |
| C13-C12 | RC-RD | C13-C12 | 1.475 | 1.43 | 1.400 | 215894.4 | 225852.32 | 392459.2 |
| C12-C11 | RD-RE | C12-C11 | 1.327 | 1.38 | 1.400 | 428441.6 | 346518.88 | 392459.2 |
| C11-C10 | RE-RF | C11-C10 | 1.475 | 1.43 | 1.400 | 215894.4 | 258403.84 | 392459.2 |
| C10-C9 | RF-RG | C10-C9 | 1.327 | 1.39 | 1.400 | 428441.6 | 315557.28 | 392459.2 |
| C9-C8 | RG-RH | C9-C8 | 1.475 | 1.45 | 1.430 | 215894.4 | 228362.72 | 360597. |
| C8-C7 | RH-RI | C8-C7 | 1.327 | 1.37 | 1.380 | 428441.6 | 364091.68 | 422500.8 |
| | RI-RJ | C7-C6 | | 1.41 | 1.440 | | 211375.68 | 348216.5 |
| | RJ-RK | C6-C5 | | 1.33 | 1.380 | | | |

**Table S2. Bond-length calculation of the RSBH+ QC system.** Only the bonds, which potentially contribute to the delocalized pi-electron system are shown. All numbers are given in Å.

| **Bond** | **RHF** | **B3LYP** | **PBE** | **MP2** |
|---|---|---|---|---|
| NZ-C15 | 1.3056 | 1.3345 | 1.3422 | 1.3260 |
| C15-C14 | 1.3909 | 1.3859 | 1.3922 | 1.3942 |
| C14-C13 | 1.3867 | 1.4101 | 1.4157 | 1.3959 |
| C13-C12 | 1.4227 | 1.4072 | 1.4125 | 1.4137 |
| C12-C11 | 1.3613 | 1.3916 | 1.3990 | 1.3831 |
| C11-C10 | 1.4249 | 1.4035 | 1.4063 | 1.4080 |
| C10-C9 | 1.3594 | 1.3952 | 1.4052 | 1.3881 |
| C9-C8 | 1.4579 | 1.4301 | 1.4302 | 1.4314 |
| C8-C7 | 1.3405 | 1.3746 | 1.3860 | 1.3729 |
| C7-C6 | 1.4740 | 1.4460 | 1.4429 | 1.4459 |
| C6-C5 | 1.3464 | 1.3798 | 1.3936 | 1.3788 |

**Table S3 Non-bonded parameters of retinal within the OPLS/AA forcefield.** Gray background are standard lysine parameters.

| atom name | Bond-type | Sigma/nm | Epsilon/(kJ/mol) | charges/e |
|---|---|---|---|---|
| CE | RCE | 3.50000e-01 | 2.76144e-01 | 0.032 |
| HE1 | HC | 2.50000e-01 | 1.25520e-01 | 0.095 |
| HE2 | HC | 2.50000e-01 | 1.25520e-01 | 0.095 |
| NZ | RNZ | 3.25000e-01 | 7.11280e-01 | -0.306 |
| HZ | H | 0.00000e+00 | 0.00000e+00 | 0.239 |
| C15 | RC15 | 3.55000e-01 | 3.17984e-01 | 0.153 |
| H15 | HC | 2.42000e-01 | 2.42000e-01 | 0.112 |
| C14 | RC14 | 3.55000e-01 | 3.17984e-01 | -0.132 |
| H14 | HC | 2.42000e-01 | 2.42000e-01 | 0.070 |
| C13 | RC13 | 3.55000e-01 | 2.92889e-01 | 0.114 |
| C12 | RC12 | 3.55000e-01 | 3.17984e-01 | -0.106 |
| H12 | HC | 2.42000e-01 | 2.42000e-01 | 0.070 |
| C11 | RC11 | 3.55000e-01 | 3.17984e-01 | 0.029 |
| H11 | HC | 2.42000e-01 | 2.42000e-01 | 0.087 |
| C10 | RC10 | 3.55000e-01 | 3.17984e-01 | -0.102 |
| H10 | HC | 2.42000e-01 | 2.42000e-01 | 0.069 |

| | | | | |
|---|---|---|---|---|
| C9 | RC09 | 3.55000e-01 | 2.92889e-01 | 0.101 |
| C8 | RC08 | 3.55000e-01 | 3.17984e-01 | -0.101 |
| H8 | HC | 2.42000e-01 | 2.42000e-01 | 0.073 |
| C7 | RC07 | 3.55000e-01 | 3.17984e-01 | 0.005 |
| H7 | HC | 2.42000e-01 | 2.42000e-01 | 0.080 |
| C6 | RC06 | 3.55000e-01 | 3.17984e-01 | -0.035 |
| C5 | RC05 | 3.55000e-01 | 3.17984e-01 | 0.069 |
| C4 | CT | 3.50000e-01 | 2.76144e-01 | -0.107 |
| H41 | HC | 2.50000e-01 | 1.25520e-01 | 0.070 |
| H42 | HC | 2.50000e-01 | 1.25520e-01 | 0.070 |
| C3 | CT | 3.50000e-01 | 2.76144e-01 | -0.096 |
| H31 | HC | 2.50000e-01 | 1.25520e-01 | 0.061 |
| H32 | HC | 2.50000e-01 | 1.25520e-01 | 0.061 |
| C2 | CT | 3.50000e-01 | 2.76144e-01 | -0.094 |
| H21 | HC | 2.50000e-01 | 1.25520e-01 | 0.055 |
| H22 | HC | 2.50000e-01 | 1.25520e-01 | 0.055 |
| C1 | CT | 3.50000e-01 | 2.76144e-01 | 0.041 |
| C16 | CT | 3.50000e-01 | 2.76144e-01 | -0.180 |
| H161 | HC | 2.50000e-01 | 1.25520e-01 | 0.060 |
| H162 | HC | 2.50000e-01 | 1.25520e-01 | 0.060 |
| H163 | HC | 2.50000e-01 | 1.25520e-01 | 0.060 |
| C17 | CT | 3.50000e-01 | 2.76144e-01 | -0.180 |
| H171 | HC | 2.50000e-01 | 1.25520e-01 | 0.060 |
| H172 | HC | 2.50000e-01 | 1.25520e-01 | 0.060 |
| H173 | HC | 2.50000e-01 | 1.25520e-01 | 0.060 |
| C18 | CT | 3.50000e-01 | 2.76144e-01 | -0.185 |
| H181 | HC | 2.50000e-01 | 1.25520e-01 | 0.075 |
| H182 | HC | 2.50000e-01 | 1.25520e-01 | 0.075 |
| H183 | HC | 2.50000e-01 | 1.25520e-01 | 0.075 |
| C19 | CT | 3.50000e-01 | 2.76144e-01 | -0.186 |
| H191 | HC | 2.50000e-01 | 1.25520e-01 | 0.083 |
| H192 | HC | 2.50000e-01 | 1.25520e-01 | 0.083 |
| H193 | HC | 2.50000e-01 | 1.25520e-01 | 0.083 |
| C20 | CT | 3.50000e-01 | 2.76144e-01 | -0.188 |
| H201 | HC | 2.50000e-01 | 1.25520e-01 | 0.086 |
| H202 | HC | 2.50000e-01 | 1.25520e-01 | 0.086 |
| H203 | HC | 2.50000e-01 | 1.25520e-01 | 0.086 |
| CD | CT | 3.50000e-01 | 2.76144e-01 | -0.120 |
| HD1 | HC | 2.50000e-01 | 1.25520e-01 | 0.060 |
| HD2 | HC | 2.50000e-01 | 1.25520e-01 | 0.060 |
| CG | CT | 3.50000e-01 | 2.76144e-01 | -0.120 |
| HG1 | HC | 2.50000e-01 | 1.25520e-01 | 0.060 |
| HG2 | HC | 2.50000e-01 | 1.25520e-01 | 0.060 |
| CB | CT | 3.50000e-01 | 2.76144e-01 | -0.120 |
| HB1 | HC | 2.50000e-01 | 1.25520e-01 | 0.060 |
| HB2 | HC | 2.50000e-01 | 1.25520e-01 | 0.060 |
| N | N | 3.25000e-01 | 7.11280e-01 | -0.500 |
| H | H | 0.00000e+00 | 0.00000e+00 | 0.300 |
| CA | CT_2 | 3.50000e-01 | 2.76144e-01 | 0.140 |
| HA | HC | 2.50000e-01 | 1.25520e-01 | 0.060 |
| C | C | 3.75000e-01 | 4.39320e-01 | 0.500 |
| O | O | 2.96000e-01 | 8.78640e-01 | -0.500 |

**Table S4 Bond parameters of retinal within the OPLS/AA forcefield.** Gray background are standard lysine parameters.

| bond (atom type-atom type) | $r_0$/ nm | force constant $k_B$/ (kJ/(mol nm$^2$)) |
|---|---|---|
| RC05-CT | 0.15100 | 265265.6 |
| RC05-RC06 | 0.13800 | 422500.8 |
| RC06-CT | 0.15100 | 265265.6 |
| RC07-RC06 | 0.14400 | 348216.5 |
| RC07-HC | 0.10800 | 284512.0 |
| RC08-RC07 | 0.13800 | 412078.5 |
| RC08-HC | 0.10800 | 284512.0 |
| RC09-RC08 | 0.14300 | 360597.2 |
| RC09-CT | 0.15100 | 265265.6 |
| RC10-RC09 | 0.14000 | 392459.2 |
| RC10-HC | 0.10800 | 284512.0 |
| RC11-RC10 | 0.14000 | 392459.2 |
| RC11-HC | 0.10800 | 284512.0 |
| RC12-RC11 | 0.14000 | 392459.2 |
| RC12-HC | 0.10800 | 284512.0 |
| RC13-RC12 | 0.14000 | 392459.2 |
| RC13-CT | 0.15100 | 265265.6 |
| RC14-RC13 | 0.14000 | 392459.2 |
| RC14-HC | 0.10800 | 284512.0 |
| RC15-RC14 | 0.14000 | 392459.2 |
| RC15-HC | 0.10800 | 284512.0 |
| RNZ-RC15 | 0.13400 | 402750.8 |
| RNZ-H | 0.10001 | 363171.2 |
| RCE-RNZ | 0.14650 | 344761.6 |
| CT-RCE | 0.15290 | 224262.4 |
| RCE-HC | 0.10800 | 284512.0 |
| RC05-RC18 | 0.15220 | 294346.1 |
| RC18-HC | 0.10800 | 284512.0 |
| C-O | 0.12290 | 476976.0 |
| C-N | 0.13350 | 410032.0 |
| C-CT_2 | 0.15220 | 265265.6 |
| CT-CT_2 | 0.15290 | 224262.4 |
| CT_2-HC | 0.10900 | 284512.0 |
| CT_2-N | 0.14490 | 282001.6 |
| CT-HC | 0.10900 | 284512.0 |
| H-N | 0.10100 | 363171.2 |
| CT-CT | 0.15290 | 224262.4 |

**Table S5 Bond angle parameters of retinal within the OPLS/AA forcefield.** Gray background are standard lysine parameters.

| Angle (atom type-atom type-atom type) | $F_0$ / ° | force constant $k_B$/ (kJ/(mol nm$^2$)) |
|---|---|---|

| | | |
|---|---|---|
| CT-CT-RC05 | 109.500 | 527.184 |
| HC-CT-RC05 | 109.500 | 292.880 |
| RC18-RC05-RC06 | 120.000 | 585.760 |
| RC18-RC05-CT | 120.000 | 585.760 |
| RC05-RC18-HC | 109.500 | 292.880 |
| HC-RC18-HC | 107.800 | 276.144 |
| CT-CT-RC06 | 120.000 | 527.184 |
| RC06-RC05-CT | 120.000 | 527.184 |
| CT-RC06-RC05 | 120.000 | 527.184 |
| RC07-RC06-RC05 | 120.000 | 527.184 |
| RC07-RC06-CT | 120.000 | 527.184 |
| RC08-RC07-RC06 | 120.000 | 527.184 |
| RC06-RC07-HC | 117.000 | 292.880 |
| RC08-RC07-HC | 117.000 | 292.880 |
| RC09-RC08-RC07 | 120.000 | 527.184 |
| RC09-RC08-HC | 117.000 | 292.880 |
| HC-RC08-RC07 | 120.000 | 292.880 |
| RC10-RC09-RC08 | 120.000 | 527.184 |
| RC08-RC09-CT | 120.000 | 585.760 |
| RC10-RC09-CT | 120.000 | 585.760 |
| RC09-CT-HC | 109.500 | 292.880 |
| RC11-RC10-RC09 | 120.000 | 527.184 |
| RC09-RC10-HC | 117.000 | 292.880 |
| RC11-RC10-HC | 117.000 | 292.880 |
| RC12-RC11-RC10 | 120.000 | 527.184 |
| RC12-RC11-HC | 117.000 | 292.880 |
| HC-RC11-RC10 | 117.000 | 292.880 |
| RC13-RC12-RC11 | 120.000 | 527.184 |
| RC13-RC12-HC | 117.000 | 292.880 |
| HC-RC12-RC11 | 117.000 | 292.880 |
| RC14-RC13-RC12 | 120.000 | 527.184 |
| RC12-RC13-CT | 120.000 | 585.760 |
| RC14-RC13-CT | 120.000 | 585.760 |
| RC13-CT-HC | 109.500 | 292.880 |
| RC15-RC14-RC13 | 120.000 | 527.184 |
| RC13-RC14-HC | 117.000 | 292.880 |
| RC15-RC14-HC | 117.000 | 292.880 |
| RNZ-RC15-RC14 | 120.000 | 527.184 |
| RNZ-RC15-HC | 117.000 | 317.984 |
| HC-RC15-RC14 | 117.000 | 292.880 |
| RCE-RNZ-RC15 | 120.000 | 527.184 |
| RC15-RNZ-H | 117.000 | 317.984 |
| H-RNZ-RCE | 117.000 | 317.984 |
| CT-RCE-RNZ | 109.500 | 669.440 |
| RNZ-RCE-HC | 109.500 | 292.880 |
| HC-RCE-HC | 107.800 | 276.144 |

| | | |
|---|---|---|
| HC-RCE-CT | 109.500 | 313.800 |
| CT-CT-RCE | 109.500 | 488.273 |
| RCE-CT-HC | 110.700 | 313.800 |
| C-N-H | 119.800 | 292.880 |
| C-N-CT_2 | 121.900 | 418.400 |
| C-CT_2-N | 110.100 | 527.184 |
| CT-CT-CT | 112.700 | 488.273 |
| HC-CT-HC | 107.800 | 276.144 |
| CT-CT-HC | 110.700 | 313.800 |
| CT-CT_2-HC | 110.700 | 313.800 |
| CT_2-CT-HC | 110.700 | 313.800 |
| CT_2-C-O | 120.400 | 669.440 |
| N-C-O | 122.900 | 669.440 |

**Table S6 Dihedral parameters of retinal within the OPLS/AA forcefield.** Gray background are standard lysine parameters.

| dihedral | C0 | C1 | C2 | C3 | C4 | C5 |
|---|---|---|---|---|---|---|
| CT-CT-CT-RC05 | 2.92880 | -1.46440 | 0.20920 | -1.67360 | 0 | 0 |
| CT-CT-CT-RC06 | 2.92880 | -1.46440 | 0.20920 | -1.67360 | 0 | 0 |
| RC06-CT-CT-HC | 0.62760 | 1.88280 | 0 | -2.51040 | 0 | 0 |
| RC05-CT-CT-HC | 0.62760 | 1.88280 | 0 | -2.51040 | 0 | 0 |
| RC06-RC05-CT-HC | -0.77822 | -2.33467 | 0 | 3.11290 | 0 | 0 |
| RC06-RC05-CT-CT | 6.32202 | -2.48530 | 0.70710 | -4.54382 | 0 | 0 |
| RC07-RC06-RC05-RC18 | 0 | 0 | 0 | 0 | 0 | 0 |
| CT-RC06-RC05-RC18 | 103.15960 | 0 | -103.15960 | 0 | 0 | 0 |
| CT-CT-RC05-RC18 | 29.28800 | -14.64400 | 2.09200 | -16.73600 | 0 | 0 |
| HC-CT-RC05-RC18 | 0 | 0 | 0 | 0 | 0 | 0 |
| RC06-RC05-RC18-HC | 0 | 0 | 0 | 0 | 0 | 0 |
| CT-RC05-RC18-HC | 0 | 0 | 0 | 0 | 0 | 0 |
| RC07-RC06-CT-CT | 2.92880 | -1.46440 | 0.20920 | -1.67360 | 0 | 0 |
| RC05-RC06-CT-CT | 2.92880 | -1.46440 | 0.20920 | -1.67360 | 0 | 0 |
| RC07-RC06-RC05-CT | 6.32202 | -2.48530 | 0.70710 | -4.54382 | 0 | 0 |
| CT-RC06-RC05-CT | 6.32202 | -2.48530 | 0.70710 | -4.54382 | 0 | 0 |
| RC08-RC07-RC06-RC05 | 28.03280 | 0 | -28.03280 | 0 | 0 | 0 |
| RC08-RC07-RC06-CT | 0 | 0 | 0 | 0 | 0 | 0 |
| HC-RC07-RC06-RC05 | 0 | 0 | 0 | 0 | 0 | 0 |
| HC-RC07-RC06-CT | 0 | 0 | 0 | 0 | 0 | 0 |
| RC09-RC08-RC07-HC | 0 | 0 | 0 | 0 | 0 | 0 |
| RC09-RC08-RC07-RC06 | 103.15960 | 0 | -103.15960 | 0 | 0 | 0 |
| HC-RC08-RC07-HC | 0 | 0 | 0 | 0 | 0 | 0 |
| HC-RC08-RC07-RC06 | 0 | 0 | 0 | 0 | 0 | 0 |
| RC10-RC09-RC08-HC | 0 | 0 | 0 | 0 | 0 | 0 |
| RC10-RC09-RC08-RC07 | 32.21680 | 0 | -32.21680 | 0 | 0 | 0 |
| CT-RC09-RC08-HC | 0 | 0 | 0 | 0 | 0 | 0 |

| | | | | | | |
|---|---|---|---|---|---|---|
| CT-RC09-RC08-RC07 | 0 | 0 | 0 | 0 | 0 | 0 |
| RC10-RC09-CT-HC | 0 | 0 | 0 | 0 | 0 | 0 |
| RC08-RC09-CT-HC | 0 | 0 | 0 | 0 | 0 | 0 |
| RC11-RC10-RC09-RC08 | 46.02400 | 0 | -46.02400 | 0 | 0 | 0 |
| RC11-RC10-RC09-CT | 0 | 0 | 0 | 0 | 0 | 0 |
| HC-RC10-RC09-RC08 | 0 | 0 | 0 | 0 | 0 | 0 |
| HC-RC10-RC09-CT | 0 | 0 | 0 | 0 | 0 | 0 |
| RC12-RC11-RC10-HC | 0.62760 | 1.88280 | 0 | -2.51040 | 0 | 0 |
| RC12-RC11-RC10-RC09 | 46.02400 | 0 | -46.02400 | 0 | 0 | 0 |
| HC-RC11-RC10-HC | 0.62760 | 1.88280 | 0 | -2.51040 | 0 | 0 |
| HC-RC11-RC10-RC09 | 0.62760 | 1.88280 | 0 | -2.51040 | 0 | 0 |
| RC13-RC12-RC11-HC | -0.77822 | -2.33467 | 0 | 3.11290 | 0 | 0 |
| RC13-RC12-RC11-RC10 | 46.02400 | 0 | -46.02400 | 0 | 0 | 0 |
| HC-RC12-RC11-HC | 0 | 0 | 0 | 0 | 0 | 0 |
| HC-RC12-RC11-RC10 | -0.77822 | -2.33467 | 0 | 3.11290 | 0 | 0 |
| RC14-RC13-RC12-HC | 0.62760 | 1.88280 | 0 | -2.51040 | 0 | 0 |
| RC14-RC13-RC12-RC11 | 46.02400 | 0 | -46.02400 | 0 | 0 | 0 |
| CT-RC13-RC12-HC | 0 | 0 | 0 | 0 | 0 | 0 |
| CT-RC13-RC12-RC11 | 0 | 0 | 0 | 0 | 0 | 0 |
| RC14-RC13-CT-HC | 0 | 0 | 0 | 0 | 0 | 0 |
| RC12-RC13-CT-HC | 0 | 0 | 0 | 0 | 0 | 0 |
| RC15-RC14-RC13-RC12 | 46.02400 | 0 | -46.02400 | 0 | 0 | 0 |
| RC15-RC14-RC13-CT | 0 | 0 | 0 | 0 | 0 | 0 |
| HC-RC14-RC13-RC12 | -0.77822 | -2.33467 | 0 | 3.11290 | 0 | 0 |
| HC-RC14-RC13-CT | 0 | 0 | 0 | 0 | 0 | 0 |
| RNZ-RC15-RC14-HC | 1.17152 | 3.51456 | 0 | -4.68608 | 0 | 0 |
| RNZ-RC15-RC14-RC13 | 46.02400 | 0 | -46.02400 | 0 | 0 | 0 |
| HC-RC15-RC14-HC | 0.62760 | 1.88280 | 0 | -2.51040 | 0 | 0 |
| HC-RC15-RC14-RC13 | 0.62760 | 1.88280 | 0 | -2.51040 | 0 | 0 |
| RCE-RNZ-RC15-HC- | 1.17152 | 3.51456 | 0 | -4.68608 | 0 | 0 |
| RCE-RNZ-RC15-RC14 | 69.87280 | 0 | -69.87280 | 0 | 0 | 0 |
| H-RNZ-RC15-HC | 0 | 0 | 0 | 0 | 0 | 0 |
| H-RNZ-RC15-RC14 | -1.26775 | 3.02085 | 1.74473 | -3.49782 | 0 | 0 |
| CT-RCE-RNZ-H | -1.26775 | 3.02085 | 1.74473 | -3.49782 | 0 | 0 |
| CT-RCE-RNZ-RC15 | -1.30753 | -9.85841 | 2.09220 | 10.94600 | -4.36784 | 2.52937 |
| HC-RCE-RNZ-RC15 | 1.17152 | 3.51456 | 0 | -4.68608 | 0 | 0 |
| HC-RCE-RNZ-H | 0.83680 | 2.51040 | 0 | -3.34720 | 0 | 0 |
| HC-CT-RCE-HC | 0.62760 | 1.88280 | 0 | -2.51040 | 0 | 0 |
| HC-CT-RCE-RNZ | -4.09614 | 5.08775 | 2.96645 | -3.95806 | 0 | 0 |
| CT-CT-RCE-RNZ | -5.38234 | 3.71591 | 0 | -7.13832 | 0 | 0.67829 |
| HC-RCE-CT-CT | 0.62760 | 1.88280 | 0 | -2.51040 | 0 | 0 |
| HC-RCE-CT-HC | 0.62760 | 1.88280 | 0 | -2.51040 | 0 | 0 |
| RCE-CT-CT-CT | -10.67460 | -1.41344 | 0 | 2.84145 | 0 | -10.55260 |
| RCE-CT-CT-HC | 0.62760 | 1.88280 | 0 | -2.51040 | 0 | 0 |
| C-N-CT_2-CT | 15.70255 | 31.75656 | -3.66936 | -43.78975 | 0 | 0 |
| C-N-CT_2-HC | 0 | 0 | 0 | 0 | 0 | 0 |

| | | | | | | |
|---|---|---|---|---|---|---|
| C-CT_2-N-C | -10.35749 | -29.58716 | -1.16734 | 41.11199 | 0 | 0 |
| C-CT_2-N-H | 0 | 0 | 0 | 0 | 0 | 0 |
| C-CT_2-CT-CT | -4.23421 | 7.22159 | 1.90790 | -4.89528 | 0 | 0 |
| CT-CT-CT-CT | 2.92880 | -1.46440 | 0.20920 | -1.67360 | 0 | 0 |
| CT-CT-CT-HC | 0.62760 | 1.88280 | 0 | -2.51040 | 0 | 0 |
| CT-CT_2-C-N | 5.00825 | -1.69870 | -0.37238 | -2.93716 | 0 | 0 |
| CT-CT_2-C-O | 0 | 0 | 0 | 0 | 0 | 0 |
| CT-CT_2-N-H | 0 | 0 | 0 | 0 | 0 | 0 |
| CT_2-N-C-O | 25.47638 | 0 | -25.47638 | 0 | 0 | 0 |
| CT_2-CT-CT-HC | 0.62760 | 1.88280 | 0 | -2.51040 | 0 | 0 |
| H-N-C-O | 20.50160 | 0 | -20.50160 | 0 | 0 | 0 |
| H-N-CT_2-HC | 0 | 0 | 0 | 0 | 0 | 0 |
| H-N-CT_2-C | 0 | 0 | 0 | 0 | 0 | 0 |
| HC-CT-CT-HC | 0.62760 | 1.88280 | 0 | -2.51040 | 0 | 0 |
| HC-CT-CT_2-N | 0.97069 | 2.91206 | 0 | -3.88275 | 0 | 0 |
| HC-CT-CT_2-HC | 0.62760 | 1.88280 | 0 | -2.51040 | 0 | 0 |
| HC-CT_2-C-N | 0 | 0 | 0 | 0 | 0 | 0 |
| HC-CT_2-C-O | 0 | 0 | 0 | 0 | 0 | 0 |
| N-C-CT_2-N | 10.36376 | -6.60654 | -10.49347 | 6.73624 | 0 | 0 |
| N-CT_2-C-O | 0 | 0 | 0 | 0 | 0 | 0 |

**Table S7. Summary of $\theta_{C13\text{-}C14}$ within the isomerization calculation.** Distribution of $\theta_{C13\text{-}C14}$ in the excited state and their corresponding populations (*trans-anti* %, *cis-anti* %, *cis-syn* %) after reintroducing the ground state parameter and subsequent simulation for $RSBH^+$ isomerization simulations at 90° and 270° directions.

| simulation | time / ns | h-bond | direction / ° | *trans-anti* / % | *cis-anti* / % | *cis-syn* / % | $\phi_{C13\text{-}C14}$ / ° | $\sigma_{C13\text{-}C14}$ / ° | direction / ° | *trans-anti* / % | *cis-anti* / % | *cis-syn* / % | $\phi_{C13\text{-}C14}$ / ° | $\sigma_{C13\text{-}C14}$ / ° |
|---|---|---|---|---|---|---|---|---|---|---|---|---|---|---|
| run1a | 500 | D253 | 90 | 0.65 | 0 | 0.35 | 97 | 15 | 270 | 0.75 | 0.25 | 0 | -92 | 8 |
| run1a | 550 | water | 90 | 1 | 0 | 0 | 97 | 8 | 270 | 1 | 0 | 0 | -88 | 8 |
| run1a | 600 | water | 90 | 0.9 | 0.1 | 0 | 93 | 8 | 270 | 0.95 | 0.05 | 0 | -97 | 7 |
| run1a | 650 | water | 90 | 1 | 0 | 0 | 102 | 6 | 270 | 0.9 | 0.1 | 0 | -93 | 9 |
| run1a | 700 | water | 90 | 1 | 0 | 0 | 107 | 10 | 270 | 1 | 0 | 0 | -105 | 8 |
| run1a | 750 | water | 90 | 0.9 | 0 | 0.1 | 100 | 10 | 270 | 1 | 0 | 0 | -89 | 8 |
| run1a | 800 | water | 90 | 1 | 0 | 0 | 107 | 9 | 270 | 1 | 0 | 0 | -99 | 9 |
| run1a | 850 | water | 90 | 1 | 0 | 0 | 110 | 7 | 270 | 0.95 | 0 | 0.05 | -100 | 9 |
| run1a | 900 | water | 90 | 0.8 | 0 | 0.2 | 102 | 14 | 270 | 0.4 | 0.6 | 0 | -87 | 9 |
| run1a | 950 | water | 90 | 1 | 0 | 0 | 108 | 6 | 270 | 1 | 0 | 0 | -102 | 9 |
| run1a | 1000 | water | 90 | 1 | 0 | 0 | 101 | 6 | 270 | 0.75 | 0.25 | 0 | -91 | 8 |
| run1b | 500 | D253 | 90 | 0.35 | 0.05 | 0.6 | 93 | 11 | 270 | 1 | 0 | 0 | -86 | 6 |
| run1b | 550 | D253 | 90 | 0 | 0.2 | 0.8 | 92 | 14 | 270 | 1 | 0 | 0 | -98 | 6 |
| run1b | 600 | water | 90 | 0.4 | 0 | 0.6 | 98 | 10 | 270 | 1 | 0 | 0 | -107 | 9 |
| run1b | 650 | water | 90 | 0.8 | 0 | 0.2 | 94 | 9 | 270 | 1 | 0 | 0 | -99 | 9 |
| run1b | 700 | water | 90 | 0.3 | 0.1 | 0.6 | 93 | 17 | 270 | 1 | 0 | 0 | -93 | 10 |
| run1b | 750 | water | 90 | 0.8 | 0 | 0.2 | 96 | 12 | 270 | 1 | 0 | 0 | -96 | 8 |
| run1b | 800 | water | 90 | 0.9 | 0 | 0.1 | 107 | 12 | 270 | 1 | 0 | 0 | -100 | 7 |
| run1b | 850 | water | 90 | 0.6 | 0.05 | 0.35 | 94 | 12 | 270 | 1 | 0 | 0 | -99 | 9 |
| run1b | 900 | water | 90 | 1 | 0 | 0 | 110 | 6 | 270 | 0.85 | 0.15 | 0 | -91 | 13 |
| run1b | 950 | water | 90 | 1 | 0 | 0 | 114 | 8 | 270 | 1 | 0 | 0 | -101 | 9 |
| run1b | 1000 | water | 90 | 0.5 | 0 | 0.5 | 99 | 20 | 270 | 1 | 0 | 0 | -97 | 11 |
| run2a | 500 | D253 | 90 | 1 | 0 | 0 | 110 | 6 | 270 | 1 | 0 | 0 | -99 | 9 |
| run2a | 550 | D253 | 90 | 0.1 | 0 | 0.9 | 90 | 10 | 270 | 1 | 0 | 0 | -99 | 8 |
| run2a | 600 | D253 | 90 | 0.35 | 0 | 0.65 | 92 | 19 | 270 | 1 | 0 | 0 | -95 | 6 |
| run2a | 650 | no | 90 | 1 | 0 | 0 | 110 | 12 | 270 | 1 | 0 | 0 | -106 | 6 |
| run2a | 700 | D253 | 90 | 0.25 | 0 | 0.75 | 97 | 13 | 270 | 1 | 0 | 0 | -96 | 8 |
| run2a | 750 | D253 | 90 | 0.6 | 0 | 0.4 | 104 | 16 | 270 | 1 | 0 | 0 | -97 | 7 |
| run2a | 800 | D253 | 90 | 0.8 | 0 | 0.2 | 109 | 12 | 270 | 1 | 0 | 0 | -99 | 8 |
| run2a | 850 | D253 | 90 | 1 | 0 | 0 | 108 | 8 | 270 | 1 | 0 | 0 | -94 | 5 |
| run2a | 900 | D253 | 90 | 0.4 | 0 | 0.6 | 94 | 20 | 270 | 1 | 0 | 0 | -98 | 8 |
| run2a | 950 | D253 | 90 | 0.35 | 0 | 0.65 | 103 | 16 | 270 | 1 | 0 | 0 | -95 | 6 |
| run2a | 1000 | D253 | 90 | 0.2 | 0 | 0.8 | 101 | 12 | 270 | 1 | 0 | 0 | -108 | 5 |
| run2b | 500 | D253 | 90 | 0.05 | 0 | 0.95 | 87 | 7 | 270 | 1 | 0 | 0 | -96 | 8 |
| run2b | 550 | D253 | 90 | 0.8 | 0 | 0.2 | 112 | 13 | 270 | 1 | 0 | 0 | -102 | 8 |
| run2b | 600 | water | 90 | 1 | 0 | 0 | 102 | 5 | 270 | 0.75 | 0.2 | 0.05 | -94 | 11 |
| run2b | 650 | no | 90 | 1 | 0 | 0 | 106 | 8 | 270 | 1 | 0 | 0 | -101 | 10 |
| run2b | 700 | water | 90 | 1 | 0 | 0 | 102 | 4 | 270 | 0.75 | 0.25 | 0 | -97 | 12 |
| run2b | 750 | water | 90 | 1 | 0 | 0 | 107 | 4 | 270 | 1 | 0 | 0 | -100 | 8 |
| run2b | 800 | E123 | 90 | 1 | 0 | 0 | 106 | 8 | 270 | 0.85 | 0 | 0.15 | -101 | 9 |
| run2b | 850 | no | 90 | 1 | 0 | 0 | 109 | 9 | 270 | 1 | 0 | 0 | -101 | 8 |
| run2b | 900 | water | 90 | 1 | 0 | 0 | 93 | 7 | 270 | 0.7 | 0.3 | 0 | -97 | 9 |
| run2b | 950 | water | 90 | 1 | 0 | 0 | 93 | 6 | 270 | 0.9 | 0.1 | 0 | -102 | 12 |
| run2b | 1000 | water | 90 | 1 | 0 | 0 | 109 | 5 | 270 | 0.75 | 0.25 | 0 | -90 | 8 |
| run3a | 500 | D253 | 90 | 0.6 | 0.05 | 0.35 | 93 | 13 | 270 | 1 | 0 | 0 | -98 | 7 |
| run3a | 550 | D253 | 90 | 0.1 | 0 | 0.9 | 82 | 13 | 270 | 1 | 0 | 0 | -96 | 5 |
| run3a | 600 | D253 | 90 | 0.05 | 0 | 0.95 | 91 | 10 | 270 | 1 | 0 | 0 | -91 | 7 |
| run3a | 650 | water | 90 | 1 | 0 | 0 | 104 | 6 | 270 | 0.9 | 0.05 | 0.05 | -93 | 8 |
| run3a | 700 | D253 | 90 | 0 | 0 | 1 | 84 | 8 | 270 | 1 | 0 | 0 | -103 | 5 |
| run3a | 750 | water | 90 | 0.95 | 0 | 0.05 | 97 | 11 | 270 | 1 | 0 | 0 | -89 | 6 |
| run3a | 800 | D253 | 90 | 0.15 | 0 | 0.85 | 92 | 7 | 270 | 1 | 0 | 0 | -98 | 7 |
| run3a | 850 | water | 90 | 1 | 0 | 0 | 108 | 6 | 270 | 0.85 | 0 | 0.15 | -98 | 11 |
| run3a | 900 | D253 | 90 | 0.2 | 0.25 | 0.55 | 93 | 8 | 270 | 1 | 0 | 0 | -94 | 7 |
| run3a | 950 | water | 90 | 1 | 0 | 0 | 110 | 7 | 270 | 0.95 | 0.05 | 0 | -96 | 7 |
| run3a | 1000 | D253 | 90 | 0.05 | 0 | 0.95 | 99 | 10 | 270 | 1 | 0 | 0 | -92 | 7 |
| run3b | 500 | D253 | 90 | 0.2 | 0 | 0.8 | 90 | 11 | 270 | 100 | 0 | 0 | -98 | 7 |
| run3b | 550 | D253 | 90 | 0.6 | 0 | 0.4 | 97 | 11 | 270 | 100 | 0 | 0 | -98 | 9 |
| run3b | 600 | D253 | 90 | 0.15 | 0 | 0.85 | 86 | 14 | 270 | 100 | 0 | 0 | -99 | 7 |
| run3b | 650 | D253 | 90 | 0 | 0 | 1 | 94 | 8 | 270 | 1 | 0 | 0 | -90 | 5 |
| run3b | 700 | water | 90 | 0.65 | 0 | 0.35 | 101 | 7 | 270 | 0.95 | 0.05 | 0 | -86 | 9 |
| run3b | 750 | D253 | 90 | 0 | 0 | 1 | 95 | 12 | 270 | 1 | 0 | 0 | -92 | 7 |
| run3b | 800 | D253 | 90 | 0.1 | 0 | 0.9 | 95 | 8 | 270 | 1 | 0 | 0 | -96 | 9 |
| run3b | 850 | D253 | 90 | 0 | 0 | 1 | 98 | 8 | 270 | 1 | 0 | 0 | -86 | 7 |
| run3b | 900 | D253 | 90 | 0 | 0 | 1 | 91 | 10 | 270 | 1 | 0 | 0 | -97 | 4 |
| run3b | 950 | D253 | 90 | 0 | 0 | 1 | 90 | 8 | 270 | 1 | 0 | 0 | -97 | 8 |
| run3b | 1000 | D253 | 90 | 0 | 0 | 1 | 85 | 10 | 270 | 1 | 0 | 0 | -95 | 7 |

**Table S8. Comparison of Bond-length calculations of the RSBH+ and RSB QC systems.** Only the bonds, which potentially contribute to the delocalized pi-electron system are shown. All numbers are given in Å.

| **Bond** | **PBE (RSBH+)** | **PBE (RSB)** |
|---|---|---|
| NZ-C15 | 1.3422 | 1.2964 |
| C15-C14 | 1.3922 | 1.4451 |
| C14-C13 | 1.4157 | 1.3789 |
| C13-C12 | 1.4125 | 1.4451 |
| C12-C11 | 1.3990 | 1.3760 |
| C11-C10 | 1.4063 | 1.4303 |
| C10-C9 | 1.4052 | 1.3844 |
| C9-C8 | 1.4302 | 1.4477 |
| C8-C7 | 1.3860 | 1.3734 |
| C7-C6 | 1.4429 | 1.4595 |
| C6-C5 | 1.3936 | 1.3804 |

**Table S9. Angles of the conjugated pi-electron-system.** Col. 2-5 are geometrical properties measured after QM-optimization with the B3LYP, MP2, PBE or RHF and 6-31G* basis-set. The gray backgrounded columns exhibit geometrical properties after MM-optimization either with the OPLS/AA or AMBER force field.

| **angle** | **B3LYP/ deg** | **MP2/ deg** | **PBE/ deg** | **RHF/ deg** | **OPLS-AA/ deg** |
|---|---|---|---|---|---|
| CE-NZ-C15 | 125.5082 | 125.1427 | 125.5602 | 125.6678 | 121.7875 |
| NZ-C15-C14 | 124.9555 | 124.0126 | 124.8030 | 124.3529 | 120.6040 |
| C15-C14-C13 | 124.4560 | 125.0973 | 124.4317 | 125.2738 | 125.0055 |
| C14-C13-C12 | 117.9070 | 117.6815 | 117.8293 | 117.6414 | 118.1348 |
| C13-C12-C11 | 127.0793 | 125.1554 | 127.1871 | 124.8297 | 124.2589 |
| C12-C11-C10 | 122.9597 | 122.5750 | 122.8751 | 123.3423 | 121.6819 |
| C11-C10-C9 | 127.1920 | 126.7401 | 127.3073 | 126.6380 | 124.3549 |
| C10-C9-C8 | 117.4380 | 117.2736 | 117.3699 | 117.4293 | 118.3858 |
| C9-C8-C7 | 124.3532 | 123.6922 | 124.3518 | 124.2708 | 123.1886 |
| C8-C7-C6 | 130.6300 | 129.9787 | 130.5442 | 130.4925 | 124.8901 |
| C7-C6-C5 | 116.6803 | 116.7790 | 116.6361 | 116.5076 | 118.4820 |
| C7-C6-C1 | 121.6303 | 121.4475 | 121.8737 | 121.2419 | 121.0548 |

**Table S10.** Charges of the carbon framework obtained by RHF/STO3G and RHF/6-31G*. Shown are Mullikan atomic charges and ESP fits, respectively.

| **atom name** | **STO3G mul/e** | **STO3G ESP/e** | **STO3G mul/e** | **STO3G ESP/e** |
|---|---|---|---|---|
| CE | -0.175 | -0.338 | -0.512 | -0.285 |
| HE1 | 0.061 | 0.091 | 0.171 | 0.088 |
| HE2 | 0.062 | 0.098 | 0.171 | 0.093 |
| HE3 | 0.061 | 0.098 | 0.171 | 0.093 |
| CZ | 0.042 | 0.037 | -0.169 | 0.037 |
| HZ | 0.052 | 0.056 | 0.179 | 0.127 |
| C15 | -0.072 | -0.086 | -0.180 | -0.222 |

| | | | | |
|---|---|---|---|---|
| H15 | 0.053 | 0.088 | 0.184 | 0.188 |
| C14 | -0.061 | -0.208 | -0.228 | -0.268 |
| H14 | 0.052 | 0.074 | 0.184 | 0.137 |
| C13 | 0.008 | 0.276 | 0.067 | 0.279 |
| C12 | -0.059 | -0.267 | -0.210 | -0.423 |
| H12 | 0.052 | 0.106 | 0.186 | 0.197 |
| C11 | -0.068 | 0.043 | -0.200 | 0.013 |
| H11 | 0.056 | 0.068 | 0.194 | 0.145 |
| C10 | -0.066 | -0.218 | -0.234 | -0.345 |
| H10 | 0.050 | 0.084 | 0.181 | 0.166 |
| C9 | 0.013 | 0.249 | 0.075 | 0.270 |
| C8 | -0.072 | -0.258 | -0.213 | -0.409 |
| H8 | 0.053 | 0.094 | 0.189 | 0.175 |
| C7 | -0.066 | 0.096 | -0.231 | 0.129 |
| H7 | 0.056 | 0.051 | 0.195 | 0.105 |
| C6 | -0.021 | -0.365 | 0.049 | -0.583 |
| C5 | 0.018 | 0.288 | 0.009 | 0.315 |
| C4 | -0.102 | -0.185 | -0.341 | -0.164 |
| H41 | 0.052 | 0.060 | 0.166 | 0.056 |
| H42 | 0.053 | 0.064 | 0.170 | 0.057 |
| C3 | -0.095 | -0.046 | -0.316 | 0.017 |
| H31 | 0.051 | 0.043 | 0.163 | 0.025 |
| H32 | 0.051 | 0.043 | 0.162 | 0.004 |
| C2 | -0.092 | -0.259 | -0.309 | -0.259 |
| H21 | 0.055 | 0.066 | 0.159 | 0.057 |
| H22 | 0.055 | 0.058 | 0.160 | 0.052 |
| C1 | 0.044 | 0.609 | -0.046 | 0.764 |
| C16 | -0.166 | -0.511 | -0.468 | -0.585 |
| H161 | 0.051 | 0.105 | 0.152 | 0.129 |
| H162 | 0.057 | 0.110 | 0.163 | 0.147 |
| H163 | 0.049 | 0.126 | 0.168 | 0.110 |
| C17 | -0.180 | -0.421 | -0.468 | -0.442 |
| H171 | 0.060 | 0.085 | 0.169 | 0.075 |
| H172 | 0.060 | 0.095 | 0.162 | 0.109 |
| H173 | 0.060 | 0.102 | 0.152 | 0.103 |
| C18 | -0.177 | -0.413 | -0.539 | -0.423 |
| H181 | 0.061 | 0.110 | 0.177 | 0.104 |
| H182 | 0.061 | 0.096 | 0.175 | 0.117 |
| H183 | 0.057 | 0.106 | 0.179 | 0.116 |
| C19 | -0.176 | -0.429 | -0.537 | -0.394 |
| H191 | 0.062 | 0.113 | 0.177 | 0.110 |
| H192 | 0.061 | 0.113 | 0.175 | 0.112 |
| H193 | 0.061 | 0.109 | 0.175 | 0.110 |
| C20 | -0.175 | -0.379 | -0.533 | -0.379 |
| H201 | 0.062 | 0.095 | 0.176 | 0.085 |
| H202 | 0.061 | 0.098 | 0.175 | 0.086 |
| H203 | 0.061 | 0.098 | 0.175 | 0.087 |

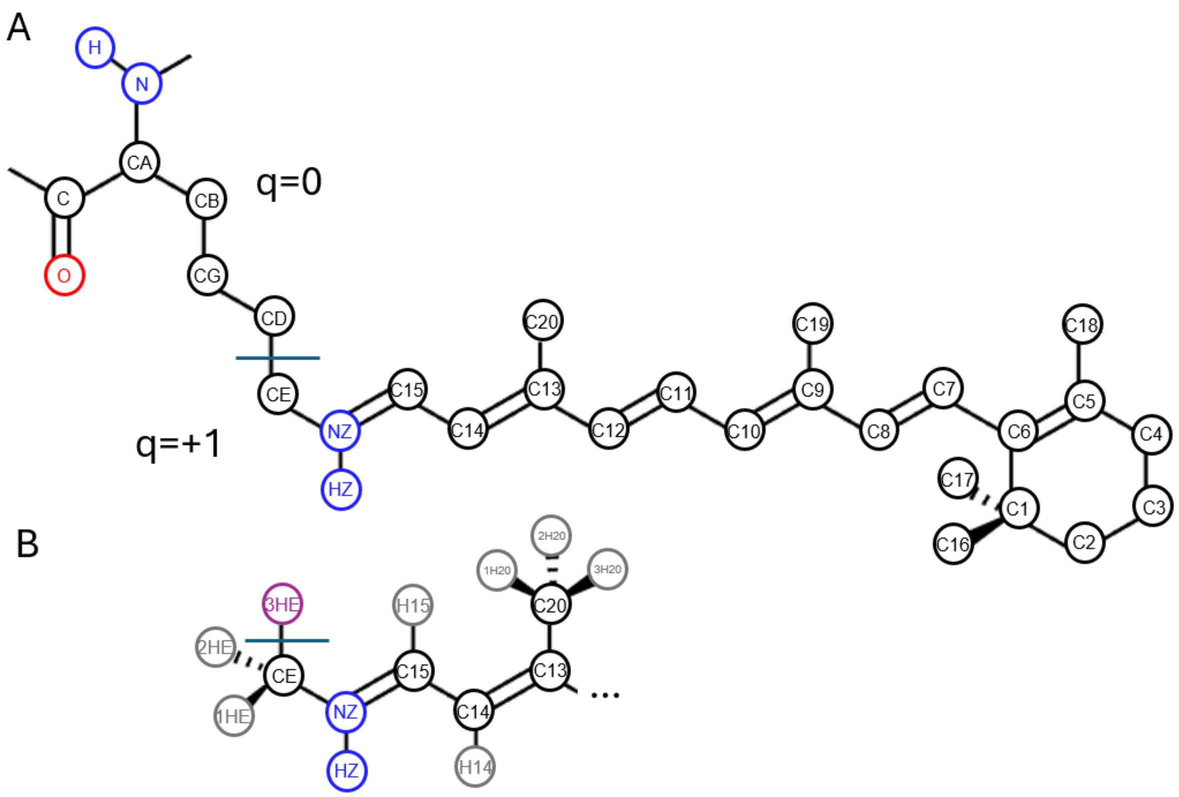


**Figure S1 Overview of the atom name and charges groups of the retinal parameters in the OPLS/AA forcefield.** A The uncharged group of the lysine core group remains unchanged, while new parameters are implemented for the charged part (QC system). B A detailed part of RSBH+ with explicit hydrogens. 3HE, marked in magenta, saturates the QC system, but is not part of the new implemented RET-Residue.

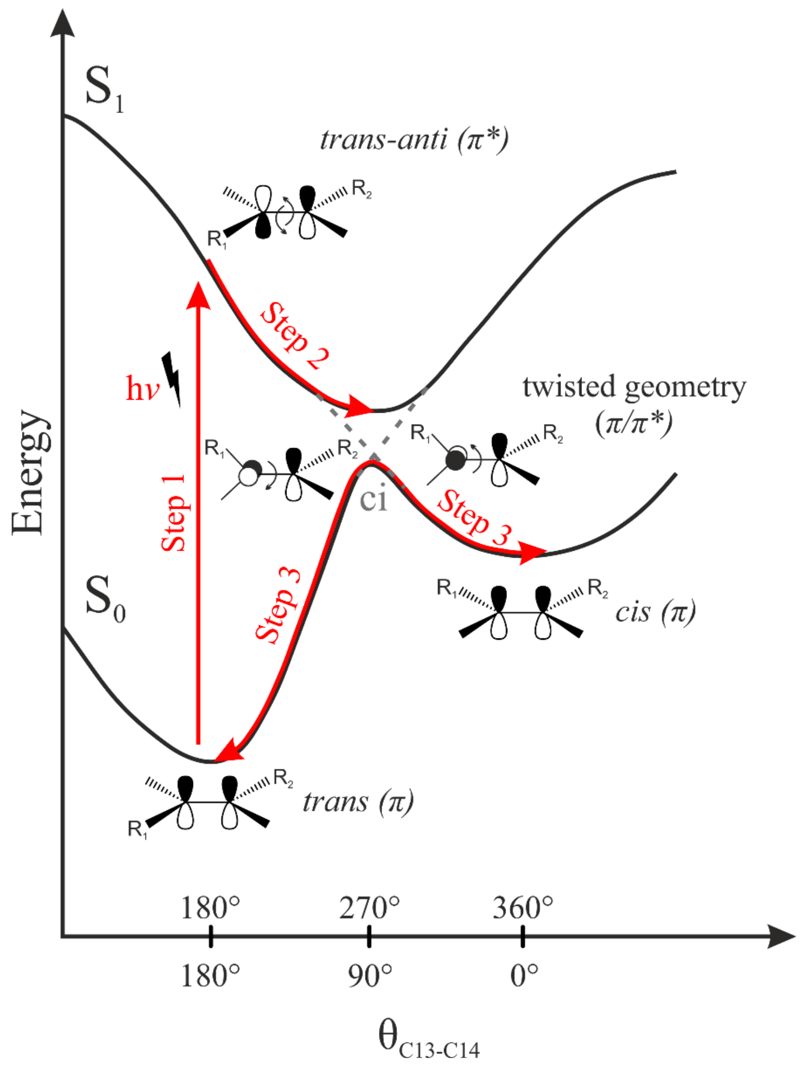


**Figure S2. Schematic illustration of the photo-isomerization process.** The photoexcitation S0 → S1 (step 1) induces a change in the bonding pattern from bonding to antibonding character between carbon atoms C13 and C14 thus favoring a 90° or 270° minimum for the passing $\theta_{C13\text{-}C14}$ torsion. The geometry is expected to equilibrate within femtoseconds around one of these new minima (step 2). At these minima geometries, the potential energy surfaces of S0 and S1 are extremely close and form an avoided conical intersection. Passing through this energy gap the molecule falls back into the ground state and the double bond character is recovered (step 3).

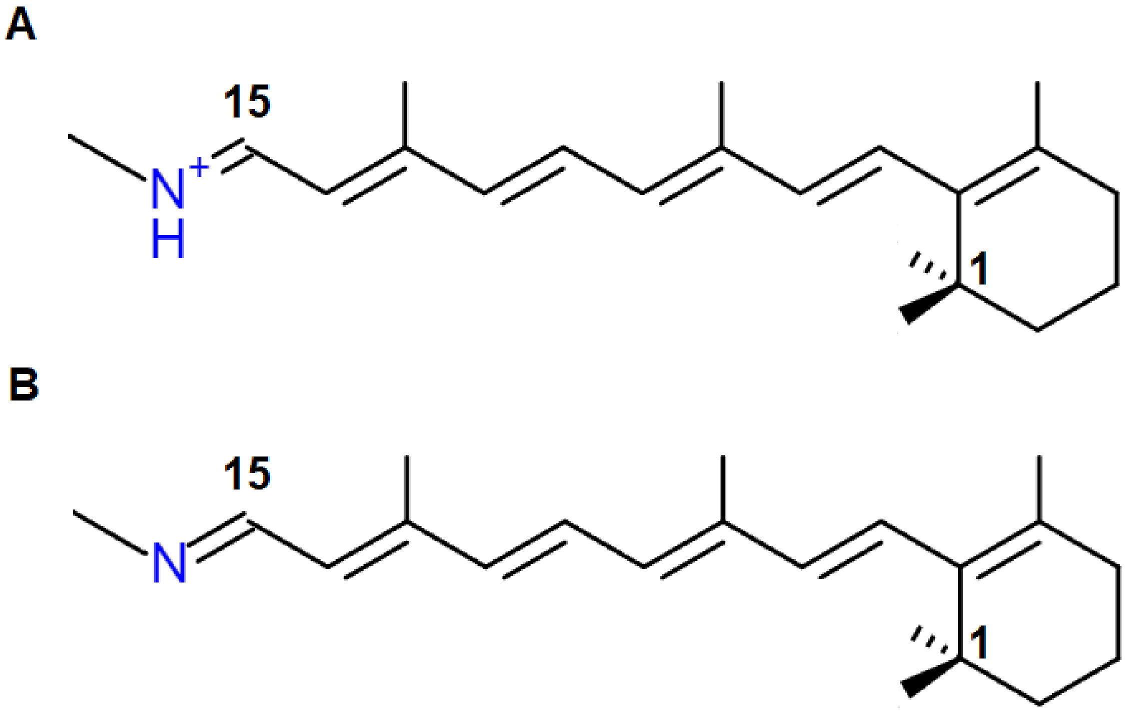


**Figure S3 QC systems. A)** the charged version of the RSBH+ QC system with protonated Schiff-base. **B)** the uncharged version of the RSB QC-system with deprotonated Schiff-base. For a better overview the skeletal formula is shown.

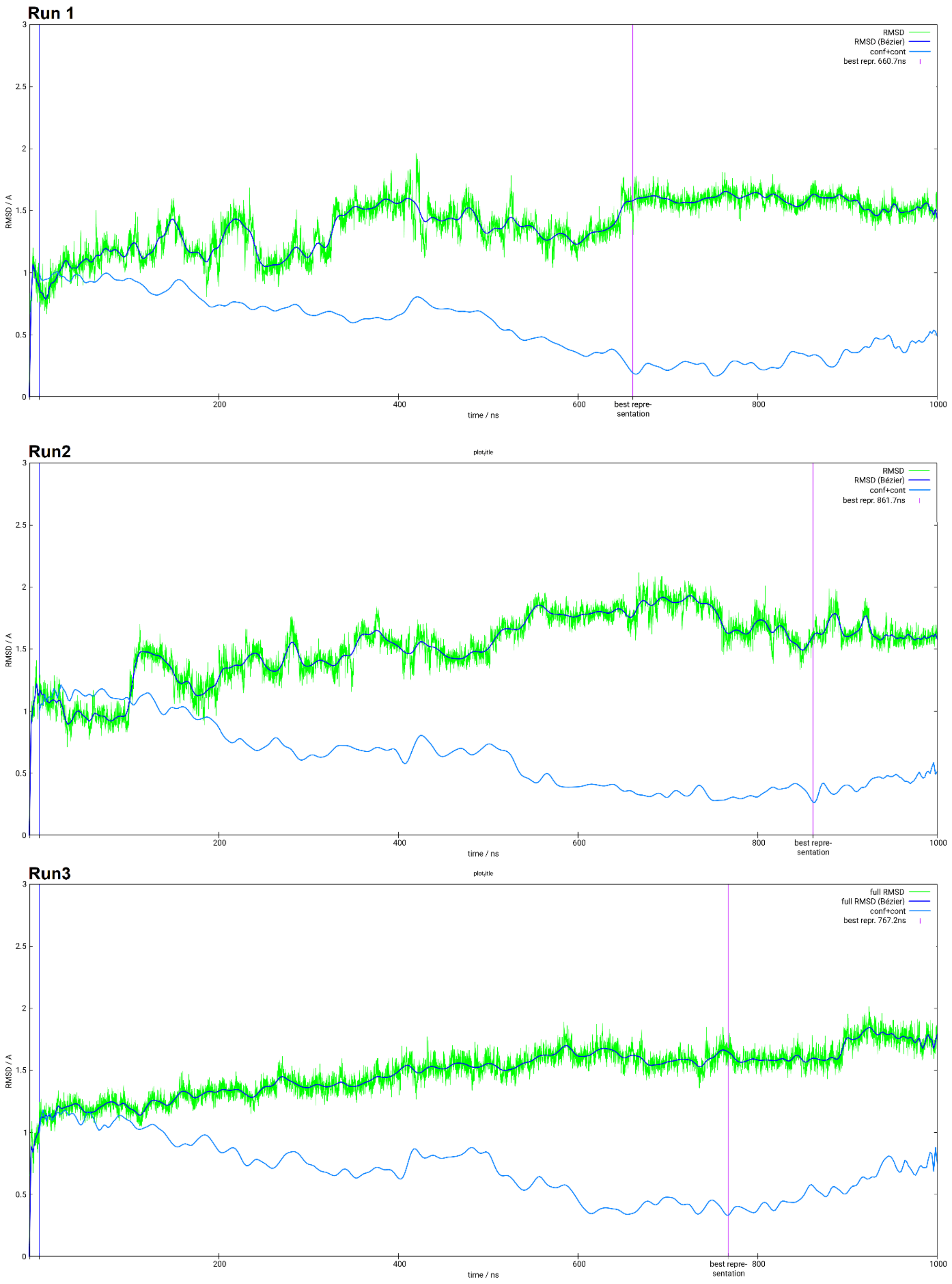


**Figure S4**. RMSD and conformation+contact values of run1,2 and 3. In green are the unfiltered values and in blue is the beziert fit. Marine shows the combined contact and conformation value. The line in magenta shows the timepoint of the representative structure.

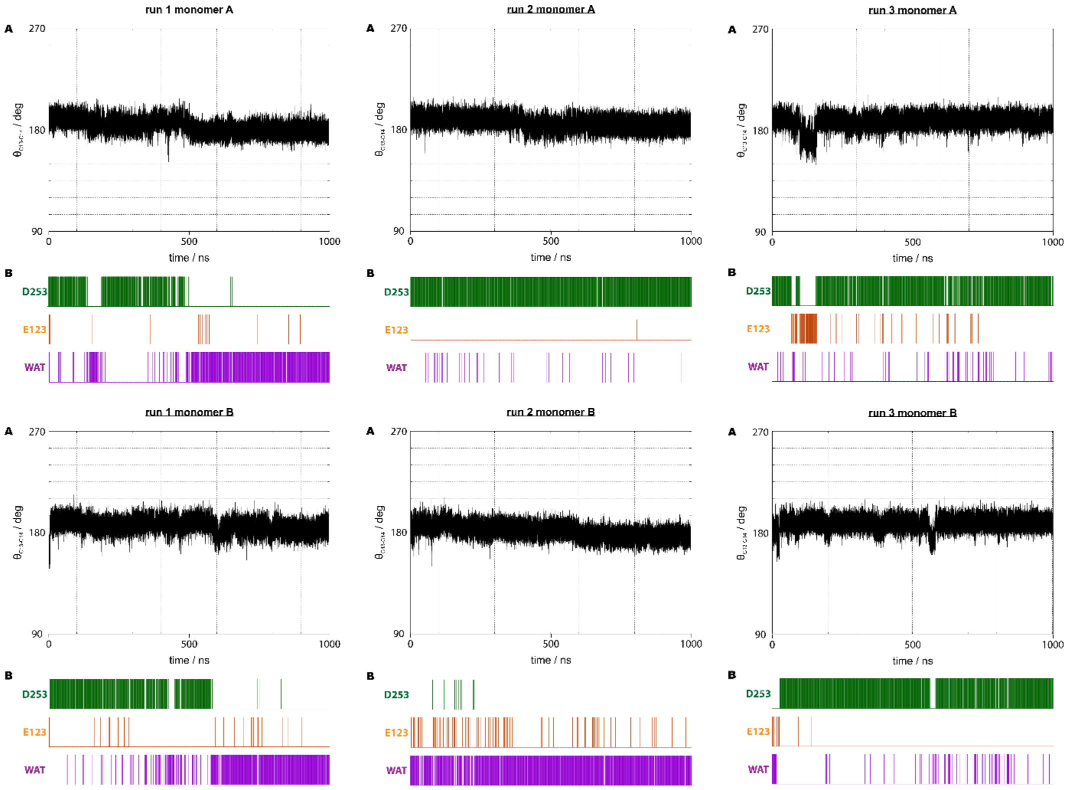


**Figure S5. Histograms of retinal configurational distribution in the ground state (S0) and in the excite state (S1).** To probe the restraint force constant impact, we compared the $\theta_{C13\text{-}C14}$ distribution for force constant values of 100 kJ/mol with excited state restrain $\theta_{C13\text{-}C14}$ to 90° (light blue) and 270° (light red), 150 kJ/mol (blue; red), and 200 kJ/mol (dark blue; dark red).

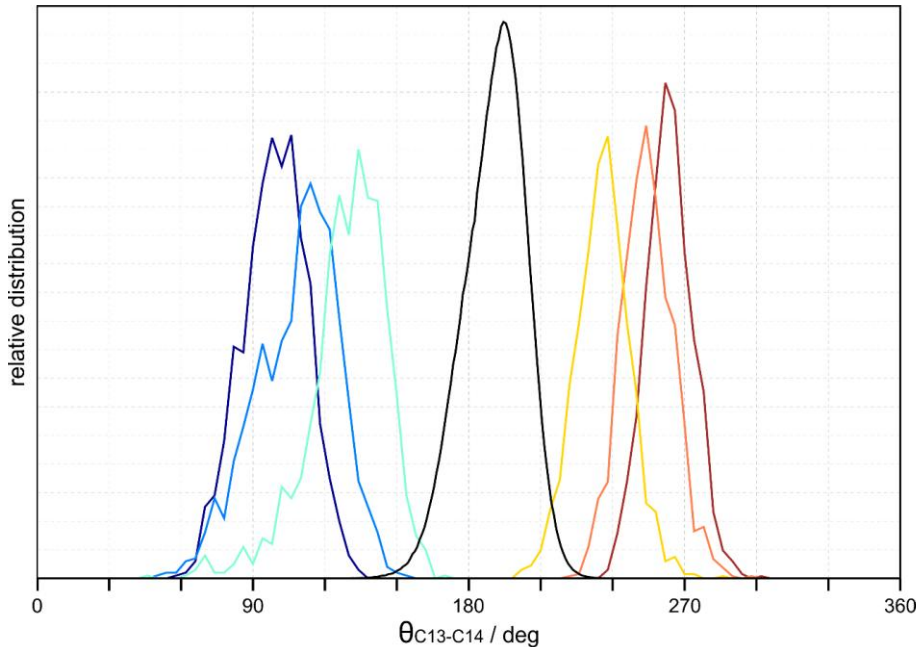


**Figure S6. Histograms of retinal configurational distribution in the ground state (S0) and in the excite state (S1).** To probe the restraint force constant impact, we compared the $\theta_{C13\text{-}C14}$ distribution for force constant values of 100 kJ/mol with excited state restrain $\theta_{C13\text{-}C14}$ to 90° (light blue) and 270° (light red), 150 kJ/mol (blue; red), and 200 kJ/mol (dark blue; dark red).

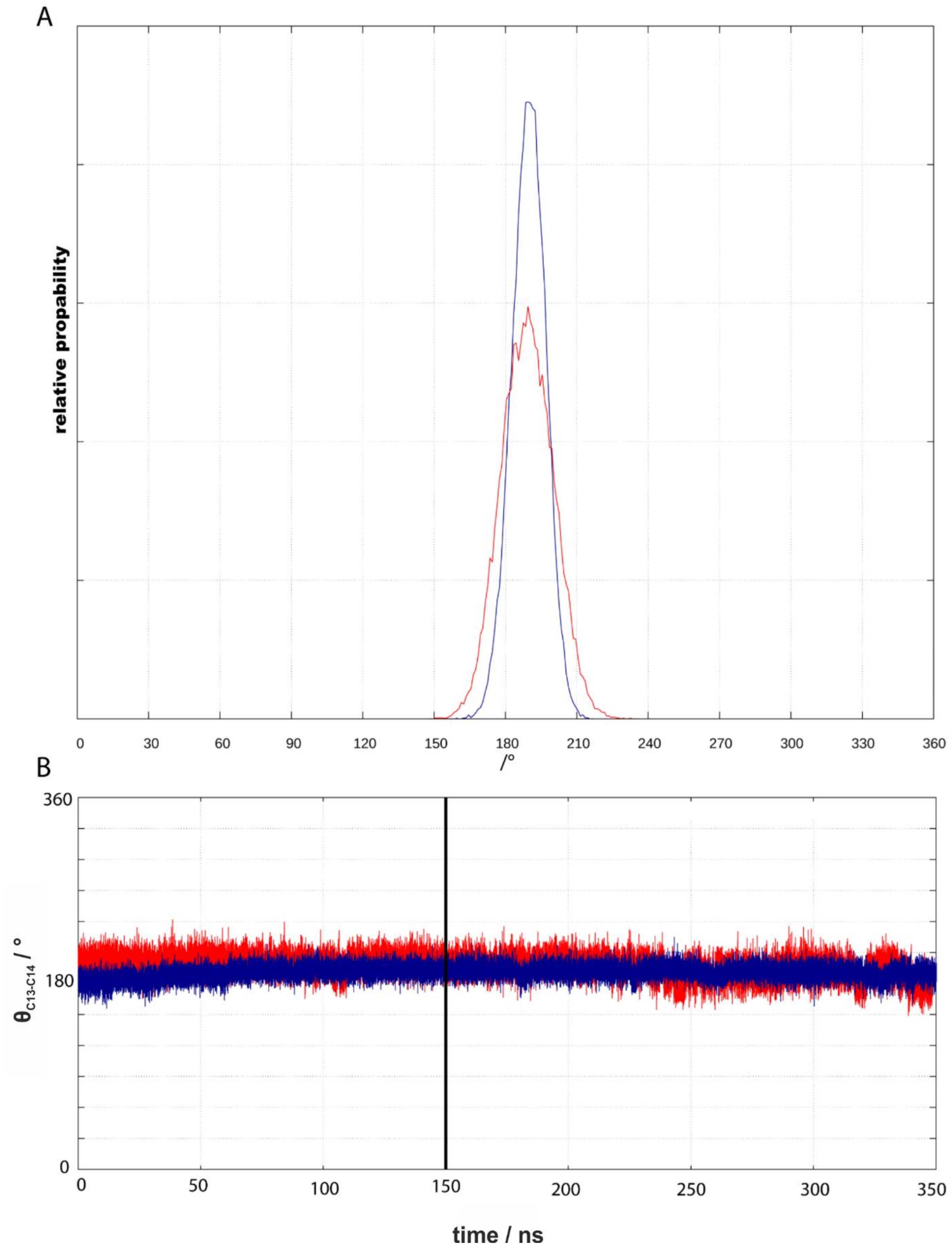


**Figure S7.** A: Relative probability of ground state geometries. Compared is the distribution of $\theta_{C13\text{-}C14}$ angle in ground-state geometries of simulations with different force constants. Only geometries from 150-350 ns of both runs are considered. Blue shows the distribution while the force constant was doubled compared to the standard parameters in red. B shows the dihedral angle of both runs depending on the simulation time. The black line indicated that both angles are equilibrated at this point, since the blue plot shows a slope before this point and runs consistently without slope afterwards.

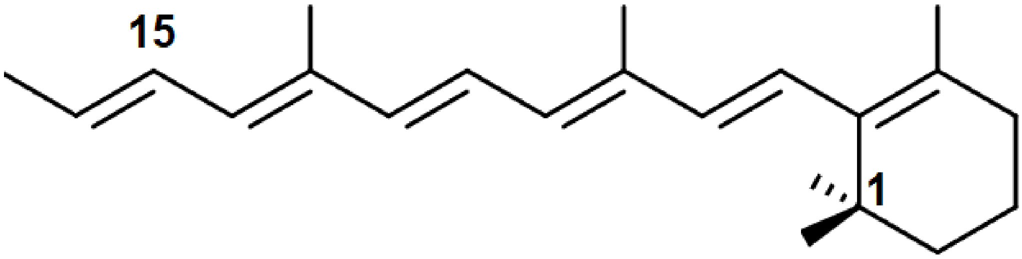


**Figure S8 QC system of the carbon framework for charge calculation**. NZ is substituted by a carbon, resulting in a net charge of 0e. For a better overview the skeletal formula is shown.

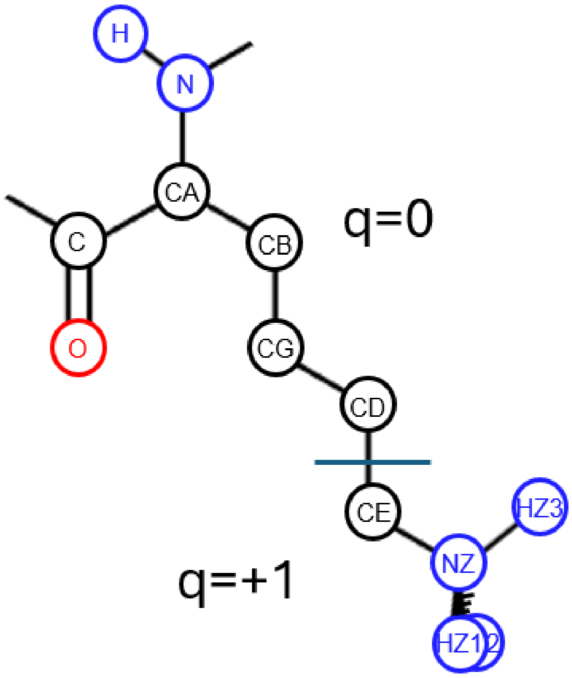


**Figure S9 Lysine charge-groups in the OPLS/AA forcefield with implicit hydrogens.** The lysine charges are separated into two groups, marked by the blue line. The charged part will be replaced by the RSBH+ QC system. Shown are the atom names.

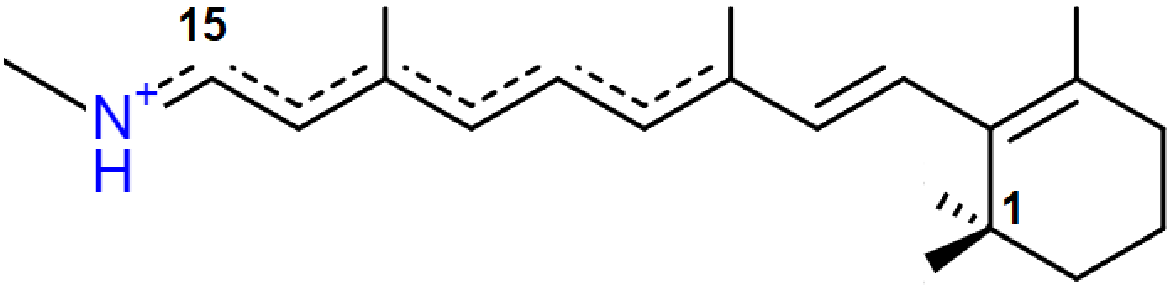


**Figure S10** QC system of RSBH+ with revised structural formula and pronounced conjugated pi electron system from NZ to the second methyl-group. After the second methyl-group the bond length alternates. For a better overview the skeletal formula is shown.

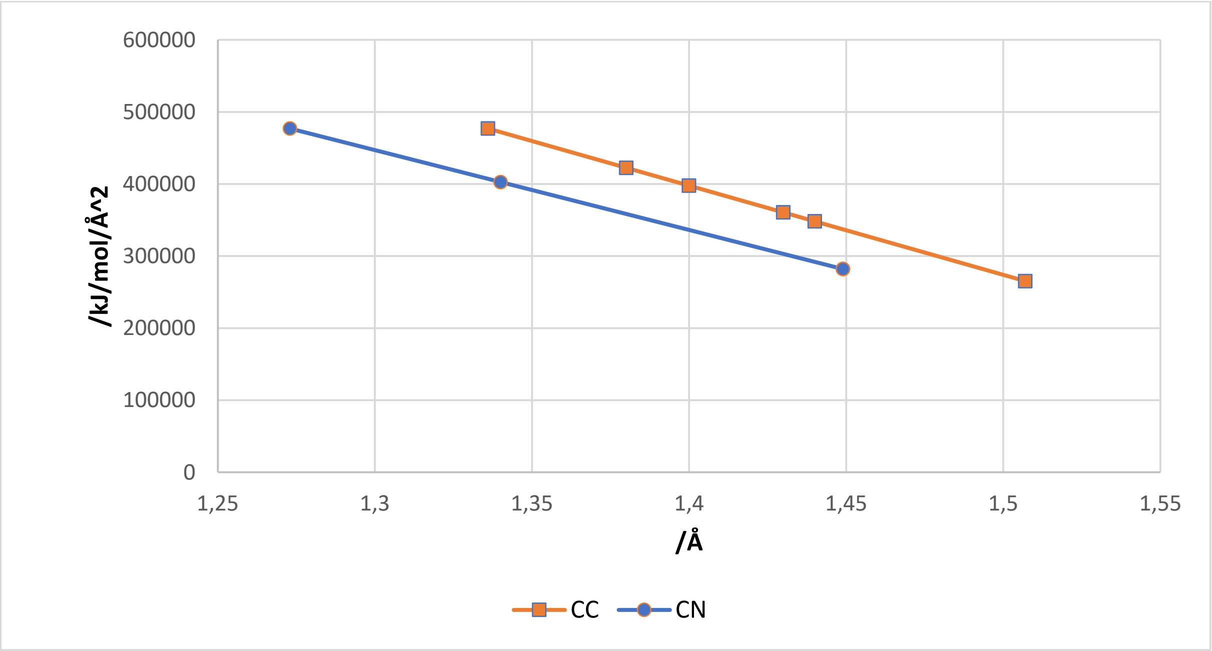


**Figure S11** Interpolation plot of CN and CC-bonds for introducing new parameters by application of the regulation for new amber-parameters converted in kJ/mol.

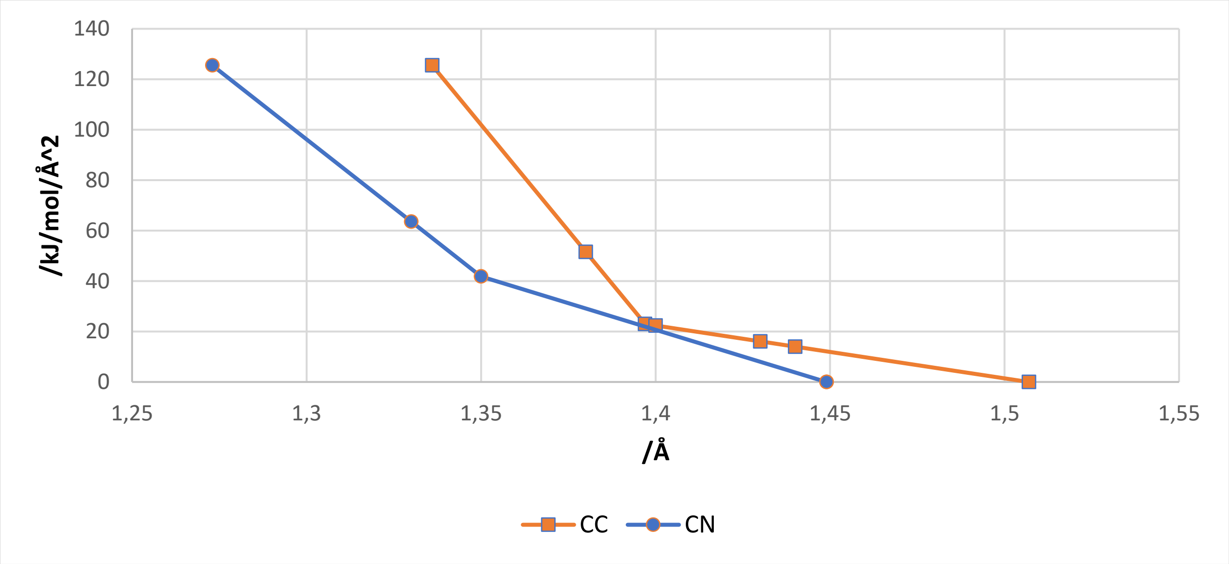


**Figure S12.** Interpolation plot of CN and CC-dihedrals for introducing new parameters by application of the regulation for new amber-parameters converted in kJ/mol.

### References

1. Jorgensen, W.L., Maxwell, D.S., Tirado-Rives, J.: Development and Testing of the OPLS All-Atom Force Field on Conformational Energetics and Properties of Organic Liquids. J. Am. Chem. Soc. **118**(45), 11225–11236 (1996). doi: 10.1021/ja9621760
2. Weiner, S.J., Kollman, P.A., Case, D.A., Singh, U.C., Ghio, C., Alagona, G., Profeta, S., Weiner, P.: A new force field for molecular mechanical simulation of nucleic acids and proteins. J. Am. Chem. Soc. **106**(3), 765–784 (1984). doi: 10.1021/ja00315a051
3. Scott J. Weiner, Peter A. Kollman, David A. Case, U. Chandra Singh, Caterina Ghio, Guliano Alagona, Salvatore Profeta, Paul Weiner: A new force field for molecular mechanical simulation of nucleic acids and proteins
4. Frisch, M.J., Trucks, G.W., Schlegel, H.B., Scuseria, G.E., Robb, M.A., Cheeseman, J.R., Scalmani, G., Barone, V., Mennucci, B., Petersson, G.A.: Gaussian 09, Revision D. 01, Gaussian, Inc., Wallingford CT. See also: URL: http://www. gaussian. com **620** (2009)
5. Pople, J.A., Nesbet, R.K.: Self-Consistent Orbitals for Radicals. The Journal of Chemical Physics **22**(3), 571–572 (1954). doi: 10.1063/1.1740120
6. Becke, A.D.: Density-functional thermochemistry. III. The role of exact exchange. The Journal of Chemical Physics **98**(7), 5648–5652 (1993). doi: 10.1063/1.464913
7. Perdew, J.P., Burke, K., Ernzerhof, M.: Generalized Gradient Approximation Made Simple. Phys. Rev. Lett. **77**(18), 3865–3868 (1996). doi: 10.1103/PhysRevLett.77.3865
8. Martin Head-Gordon, John A. Pople, Michael J. Frisch: MP2 energy evaluation by direct methods. Chemical Physics Letters **153**(6), 503–506 (1988). doi: 10.1016/0009-2614(88)85250-3
